# Synthetic biology-guided modular engineering of the secreted cutinase *Fo*Cut5a with a carbohydrate-binding module improves poly(ester-urethane) depolymerization in *Yarrowia lipolytica*

**DOI:** 10.64898/2026.10.15.759203

**Authors:** Eugenia Messina, Markella Papi, Serena Barile, Rosa De Rosa, Pasquale Scarcia, Antonino Biundo, Isabella Pisano, Evangelos Topakas, Gennaro Agrimi

## Abstract

Poly(ester-urethane) bioconversion requires efficient polymer-degrading enzymes and microbial hosts capable of processing the resulting monomers. However, the optimal modular design of secreted cutinases for depolymerizing insoluble polyester-polyurethanes remains largely unexplored. Here, we established a combinatorial synthetic biology strategy in *Yarrowia lipolytica* based on the fungal cutinase *FoCut5a* from *Fusarium oxysporum*. A library of 96 Golden Gate-assembled constructs, combining four secretion signals, four interdomain linkers, and six substrate-binding domains, was integrated site-specifically into the genome and screened for depolymerization of the polyester-polyurethane dispersion Impranil DLN-SD. The best-performing variant fused *Fo*Cut5a to the carbohydrate-binding module *Tr*CBM1. LC-MS analysis demonstrated release of adipic acid, 1,6-hexanediol, and neopentyl glycol, confirming ester-bond hydrolysis within the polyester domains of the polymer and additionally detected 6-hydroxycaproate, indicating oxidation of polyurethane-derived 1,6-hexanediol by *Y. lipolytica*. Purified *Fo*Cut5a-*Tr*CBM1 displayed unchanged activity on *p*-nitrophenyl butyrate but reduced Impranil turbidity 5.6-fold faster and increased release of 1,6-hexanediol and adipic acid by 76-fold and 220-fold, respectively, compared to the unfused enzyme. These results highlight modular enzyme engineering as an effective strategy to enhance polyester-polyurethane deconstruction and support *Y. lipolytica* as a platform for polymer bioconversion and upcycling.

## 1. Introduction

Polyurethanes (PUs) are among the most widely used synthetic polymers, with global production exceeding 20 million tonnes in 2022 and applications spanning construction materials, automotive components, textile coatings, medical devices, and adhesives (Raczyńska et al., 2024). Unlike polyolefins and polyethylene terephthalate (PET), which benefit from established mechanical and chemical recycling technologies, PUs waste remains difficult to recycle and is still largely managed through landfilling or incineration. Consequently, increasing attention has been directed toward enzymatic depolymerization as a sustainable strategy for recovering valuable PU-derived monomers and intermediates (Raczyńska et al., 2024 ; Cha et al., 2026).

PUs are often synthesized through the polyaddition of diisocyanates and polyols, forming characteristic urethane (carbamate) linkages. Depending on the polyol employed, they are classified as poly(ester-urethane) (PEU) or poly(ether-urethane) (PEUR), which differ substantially in their chemical and physical properties. Owing to the presence of hydrolysable ester bonds within their soft segments, PEU are particularly susceptible to enzymatic attack and have therefore emerged as attractive model substrates for biodegradation studies. Among the model substrates used to investigate polyurethane biodegradation, Impranil DLN-SD is the most widely adopted. This aqueous colloidal dispersion of this aliphatic PEU, whose main constituents include adipic acid, 1,6-hexanediol, and neopentyl glycol, was originally developed for textile and leather coating applications (Biffinger et al., 2015). Its colloidal structure provides a large accessible surface area for enzymatic attack, making it a valuable model system for evaluating polyurethane-degrading enzymes and elucidating depolymerization mechanisms. Accordingly, Impranil DLN-SD has become a reference substrate in both microbial and enzymatic studies of polyurethane degradation (Biffinger et al., 2015; Raczyńska et al., 2024).

The growing interest in biological PUs recycling has led to the identification of numerous microorganisms capable of degrading Impranil DLN-SD, including the filamentous fungi *Cladosporium pseudocladosporioides* T1.PL.1 (Álvarez-Barragán et al., 2016), *Cladosporium* sp. *P7* (Liu et al., 2023), among others. The degradation process is generally mediated by extracellular hydrolases, primarily esterases, lipases, and cutinases, which cleave the ester bonds present within the polyester soft segments of the polymer (Raczyńska et al., 2024;Liu et al., 2023). Among these enzymes, cutinases (EC 3.1.1.74) have attracted particular interest because of their broad substrate specificity and ability to act on insoluble polymeric materials. These serine hydrolases possess a canonical α/β-hydrolase fold and a catalytic Ser-His-Asp triad. Although originally identified as enzymes involved in plant cutin degradation, they are now recognized as efficient biocatalysts for the hydrolysis of various synthetic polyesters, including poly(ethylene terephthalate) (PET) and poly(ester-urethane), owing to their exposed active site and capacity to accommodate bulky polymeric substrates (Roman et al., 2024; Raczyńska et al., 2026). Among fungal cutinases, *Fo*Cut5a from *Fusarium oxysporum* has demonstrated activity toward both aliphatic and aromatic polyesters, including poly(ε-caprolactone) and PET-related substrates (Dimarogona et al., 2015). Furthermore, the cutinase *Pl*Cut1 from *Papiliotrema laurentii*, which belongs to the same phylogenetic clade as *Fo*Cut5a, was recently shown to hydrolyze Impranil DLN-SD and outperform a commercial lipase under the same assay conditions, highlighting the potential of fungal cutinases as poly(ester-urethane)-degrading biocatalysts (Roman et al., 2024). A close homologue of *Fo*Cut5a (99.6% amino acid identity) was recently expressed in *Aspergillus niger* and shown, in purified form, to degrade the poly(ester-urethane) dispersion Impranil DLN (Verschoor et al., 2025). These findings indicate that members of this cutinase clade are active on polyester-polyurethanes; however, the activity of *Fo*Cut5a itself on this substrate, as well as its production and secretion in a yeast host, had not previously been demonstrated.

Despite their catalytic potential, the degradation of insoluble polymers is often limited by inefficient enzyme adsorption to the substrate surface. The importance of this step is exemplified by the hydrophobic surface-binding protein HsbA, which mediates the attachment of polyurethane-degrading enzymes to the Impranil DLN-SD surface in *Cladosporium halotolerans* 6UPA1 (Zhang et al., 2022). Similarly, Gamerith et al., (2016) demonstrated that fusion of a polymer-binding module (PBM) to a polyurethane-active polyamidase increased activity on insoluble PU substrates by up to 4-fold, despite reducing activity on a soluble model substrate. These findings indicate that substrate binding and catalytic turnover can be independently optimized. Consistent with this concept, fusion of carbohydrate-binding modules (CBMs), including CBM1 from *Trichoderma reesei*, has been shown to enhance substrate interaction and hydrolytic performance of polyester-degrading cutinases on PET and related materials (Ribitsch et al., 2013;Espino-Rammer et al., 2013). Together, these studies suggest that accessory substrate binding domains (SBDs) may represent a broadly applicable strategy for improving enzymatic depolymerization of synthetic polyesters and polyurethanes. The performance of these chimeras is determined by the combined effects of the substrate-binding domain, the interdomain linker, and the secretion signal. As interactions among these modules are difficult to predict a priori, systematic exploration of the design space requires combinatorial rather than one-by-one construct assembly.

The non-conventional yeast *Yarrowia lipolytica* represents an attractive platform for the production of heterologous enzymes because of its efficient secretory machinery, genetic accessibility, and broad metabolic versatility (Fickers et al., 2005) . In recent years, the development of advanced synthetic biology and genome-engineering toolkits has greatly expanded the possibilities for the rapid and precise genetic manipulation of this organism. In particular, standardized modular toolkits for *Y. lipolytica* combine one-pot Golden Gate assembly with CRISPR/Cas9-mediated integration at defined genomic loci, enabling the construction and screening of combinatorial enzyme libraries rather than individual designs (Yuzbashev et al., 2023).

Beyond its value as a protein-production host, *Y. lipolytica* can also assimilate and convert products generated during plastic depolymerization. Our group recently reported high-titer glycolic acid production (48.4 g/L) by *Y. lipolytica* through the co-utilization of ethylene glycol, a major PET-derived monomer, and acetate, achieving a glycolic acid yield of ∼0.8 mol/mol based on consumed ethylene glycol and highlighting the potential of this yeast as a microbial chassis for integrated plastic upcycling (Messina et al., 2026).

In this study, we applied a combinatorial synthetic biology approach to engineer *Y. lipolytica* for poly(ester-urethane) depolymerization through the heterologous expression and secretion of the fungal cutinase *Fo*Cut5a, representing its first reported application for poly(ester-urethane) degradation in a yeast-based expression system. A combinatorial library of 96 expression constructs, generated by one-pot Golden Gate assembly of four signal peptides, four interdomain linkers, and six substrate-binding domains flanking the *Fo*Cut5a catalytic core, was integrated into a defined genomic locus and screened on Impranil DLN-SD to identify architectures associated with enhanced polymer degradation. Selected strains and purified enzyme variants were subsequently characterized to assess ester bond hydrolysis within the polyester domains of Impranil DLN-SD and to evaluate the impact of the substrate-binding domains on enzymatic performance. To our knowledge, this is the first report of a combinatorial approach for engineering poly(ester-urethane)-active enzymes in a yeast host, combining library generation with direct phenotypic screening on a polyurethane-containing medium.

## 2. Experimental Procedures

### 2.1 Strains and culture media

*Escherichia coli* DH5α used for plasmid construction, propagation and amplification, was cultivated in Luria Bertani (LB) medium at 37 °C with shaking (200 rpm). This medium was supplemented with 50 μg/mL spectinomycin (Sigma-Aldrich, St. Louis, MO, USA), to allow selective growth. All *Y. lipolytica* strains constructed in this study are derived from Y-4973, an uracil auxotroph and Ku70-mutant (Δ*ura3* Δ*ku70*) obtained from the W29 strain (MatA, wild-type), that shows an increased rate of homologous recombination (Yuzbashev et al., 2023). *Y. lipolytica* strains were cultivated in different liquid media at 28 °C and with orbital shaking at 200 rpm. The rich medium, used for seed culture, was YPD medium composed of 10 g/L yeast extract (Biokar Diagnostics, Allonne, France), 20 g/L peptone (Biokar Diagnostics, Allonne, France), and 20 g/L glucose (Sigma-Aldrich, St. Louis, MO, USA). Whenever required, the medium was supplemented with 350 μg/mL nourseothricin (Jena Bioscience, Jena, Germany) for the overnight growth of *Y. lipolytica* strains transformed with Cas9 expressing plasmids. The minimal medium was composed of 1.7 g/L YNB (Yeast Nitrogen Base without nitrogen source, Difco™, Becton Dickinson, Sparks, MD, USA) supplemented with 5 g/L NH_4_Cl and 50 mM phosphate buffer pH 6.8 containing potassium dihydrogen phosphate and sodium phosphate dibasic dihydrate. The dispersion of Impranil DLN-SD (Covestro Deutschland AG, Germany) was prepared starting from a pure mixture of this poly(ester-urethane) and mixing it with distilled and sterile water to obtain a 50% (v/v) concentration. Solid media were prepared by adding 20 g/L agar to the liquid media. All cultures were maintained at -80 °C in glycerol stocks.

### 2.2 Plasmid libraries and recombinant strains construction

For the construction of the esterase libraries, four esterases (*Fo*Cut5a, *Da*PUase, Se1JFR, and *Apar*EST) were each combinatorially assembled with four secretory signal peptide (SigPep) variants, generating four independent libraries comprising four plasmids. For the construction of the 96-plasmid library, four SigPep variants, four interdomain linkers, and six SBDs were combinatorially assembled with the catalytic domain of *Fusarium oxysporum* cutinase *Fo*Cut5a, generating 96 possible construct combinations. For the construction of the control library, the four SigPep variants were fused to *Fo*Cut5a in which a stop codon had been introduced at the C-terminus of the cutinase coding sequence by PCR using the primers listed in Table S1. A description of all modules is provided in Table 1. The four SigPep sequences were ordered as pairs of complementary oligonucleotides (Table S1), phosphorylated, annealed as previously described (Larroude et al., 2018), and used as double-stranded DNA fragments for assembly. The four linker sequences were obtained as a single *Y. lipolytica* codon-optimised synthetic DNA fragment (Twist Bioscience, USA) and individually amplified by PCR using specific primers (Table S1). The four esterases (*Fo*Cut5a, *Da*PUase, Se1JFR, and *Apar*EST) and six SBDs were obtained as *Y. lipolytica* codon-optimized synthetic DNA fragments (Twist Bioscience, USA). All modules were flanked by appropriately positioned BsaI recognition sites and custom four-base overhangs designed to ensure seamless and directional assembly of the constructs in the desired order during Golden Gate Assembly (GGA).

**Table 1.**
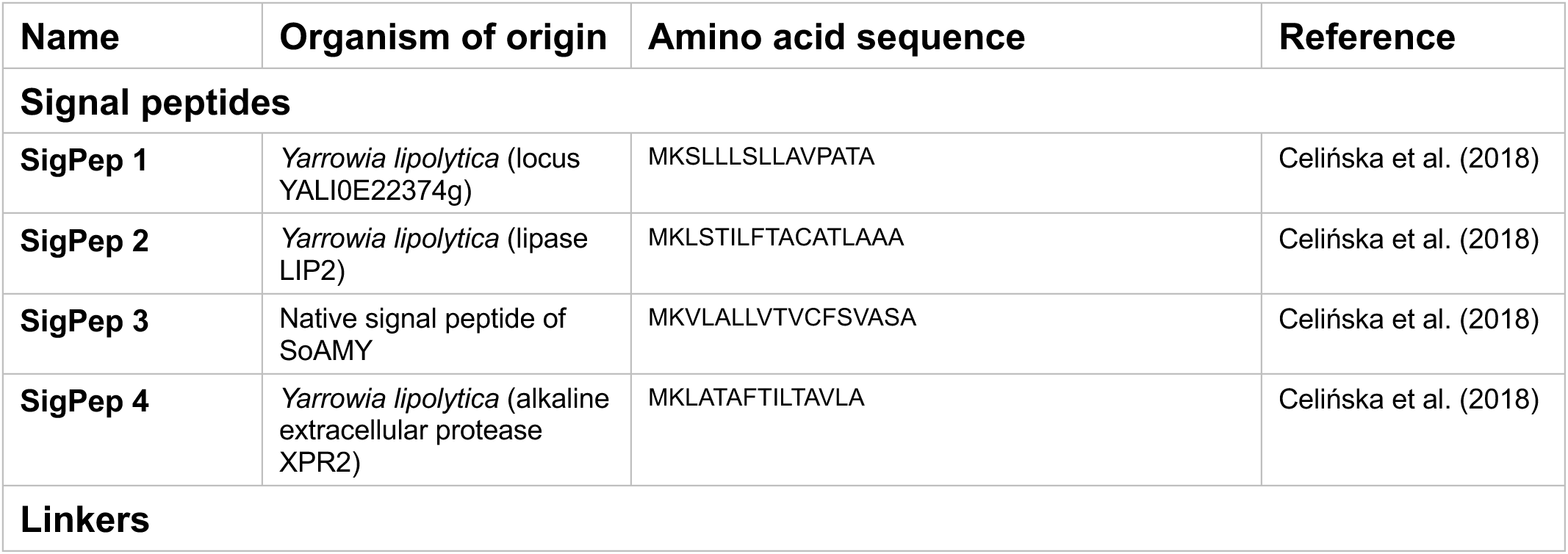
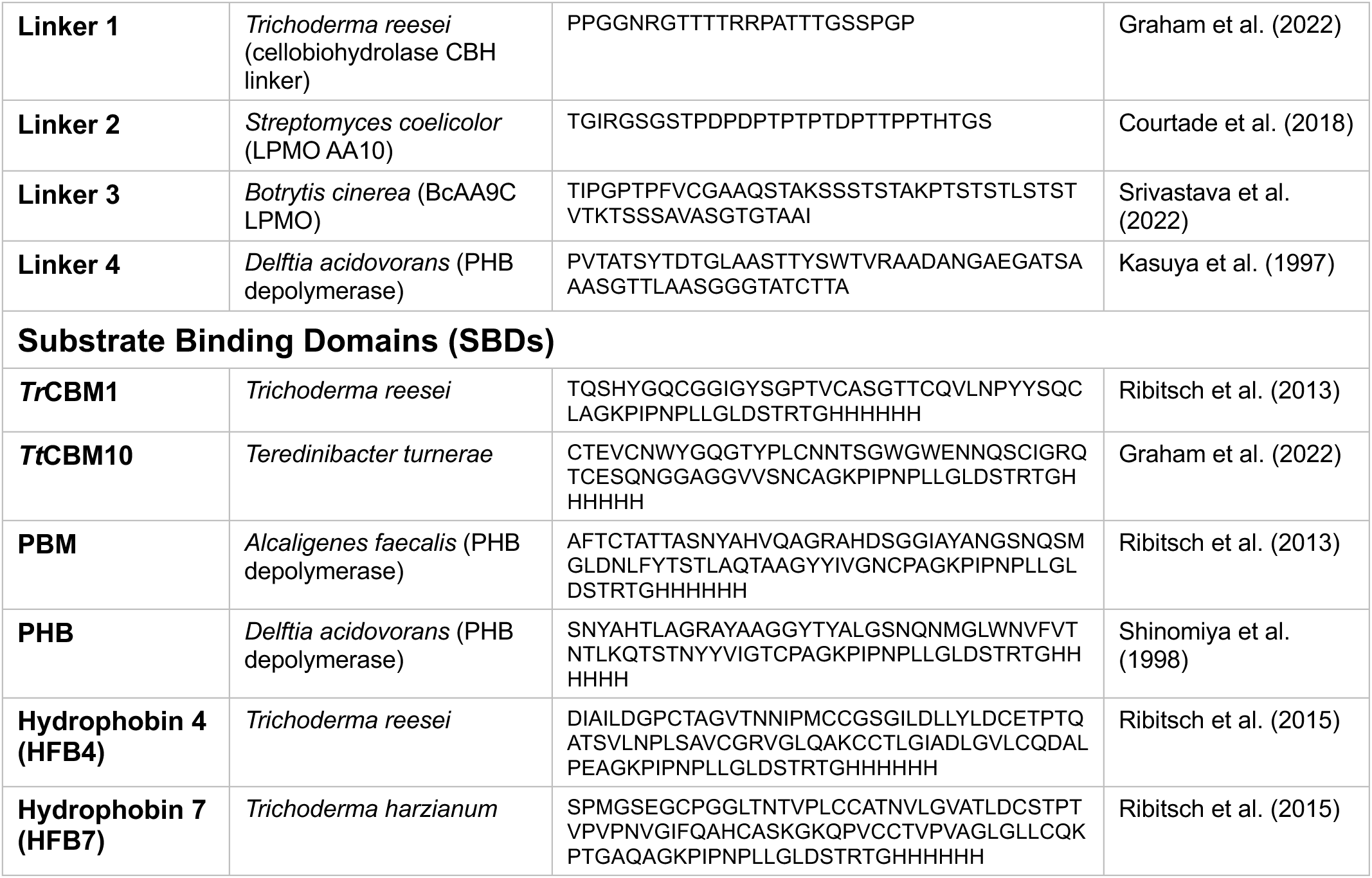
Modular components used to construct the FoCut5a fusion library. The table lists the four signal peptides, four interdomain linkers, and six substrate-binding domains (SBDs) combined in this study, together with their source organisms and amino acid sequences. The catalytic module, FoCut5a, was common to all 96 possible module combinations generated in the library. Signal peptides were included to direct protein secretion in Y. lipolytica, whereas linkers and SBDs were selected to modulate domain spacing and substrate-binding properties, respectively. All SBDs were fused to a common C-terminal V5-His₆ tag (GKPIPNPLLGLDSTRTGHHHHHH). Amino acid sequences are reported using the standard one-letter code.

The three libraries were assembled by GGA (Weber et al., 2011) using components of the *YaliCraft*® toolkit (Yuzbashev et al., 2023) . Assemblies included the Level 1 destination vector p1553, carrying the *URA3* marker and directing integration at the IntF11 locus, the Level 0 plasmids of TEF1 promoter (p1084) and the LIP2 terminator (p1595) (Table S2), and the corresponding variable modules. The assembled libraries were propagated in *E. coli* DH5α and verified by colony PCR and Sanger sequencing (Table S1). For each of the four esterase libraries, three colonies were screened by colony PCR and verified by sequencing. For the 96-member combinatorial library, 14 colonies were screened by colony PCR, of which six yielded the expected amplicon and were subsequently confirmed by sequencing. For the control library, three colonies were analyzed by colony PCR and sequencing. All tested constructs showed correct assembly. All primers used are listed in Table S1 of the Supplementary Material. Overexpression cassettes were excized by NotI digestion and co-transformed into *Y. lipolytica* together with the Cas9-helper plasmid p1135 (Table S2), targeting integration at the F11 locus. The *URA3* gene was used as the selection marker for restoration of uracil prototrophy. Transformation was performed according to Chen et al. (1997), resulting in the random integration of a single construct from either one of the four esterase libraries, the combinatorial library comprising 96 possible combinations, or the control library comprising four SigPep-*Fo*Cut5a constructs. Transformants were selected on YNB agar lacking uracil and supplemented with 1% (v/v) glucose or Impranil DLN-SD. Plates were incubated at 28 °C for up to 240 h, and hydrolysis halo formation was monitored every 24 h. Selection in the absence of uracil ensured the growth of only those strains carrying an integrated *URA3*-containing construct. Correct chromosomal integration at the F11 locus was verified by PCR using primers annealing to the genomic regions flanking the integration site (Table S1). After confirmation of cassette integration, PCR products were purified and subjected to Sanger sequencing using specific primers (Table S1). Sequence analysis enabled the identification of the individual genetic elements present in each expression cassette and, consequently, the specific construct combination integrated into the genome.

### 2.3 Shake flask cultures

Seed culture from YPD plates was grown overnight in 10 mL of YPD medium at 28 °C and 200 rpm. Pre-cultures were centrifuged, washed with sterile distilled water, and used for inoculation at initial optical density (OD_600nm_) equal to 1 into 250-mL baffled flasks containing 50 mL minimal medium composed of 1.7 g/L YNB supplemented with 5 g/L NH_4_Cl, 50 mM phosphate buffer pH 6.8 and 1% (v/v) Impranil DLN-SD dispersion. Cultivations were performed for up to 144 h at 28 °C on an orbital shaker operating at 200 rpm. Samples were collected at regular time intervals for LC-MS and HPLC analysis. All measurements were performed using at least two biological replicates, and average values with standard errors were calculated.

### 2.4 Analytical Methods

LC-MS analysis was performed to qualitatively and quantitatively assess the monomers released during Impranil DLN-SD degradation by both the engineered strains expressing the different *Fo*Cut5a cutinase variants and the corresponding purified enzymes. The supernatants were centrifuged through 0.45 µm nylon microcentrifuge filters at 13,000 rpm for 10 min at room temperature to remove any residual polymer precipitates and, when necessary, diluted prior to analysis. Metabolites were separated on a UHPLC Vanquish system using a Kinetex C18 column (150 mm length x 2.1 mm internal diameter, 2.6 µm particle size; Phenomenex) kept at 40° C. The mobile phase consisted of (A) LC-MS grade water with 0.1% formic acid and (B) Acetonitrile with 0.1% formic acid. The gradient program was as follows: 0 min, 5% B; 0-10 min, increase to 100% B; 10-15 min, 100% B; 15-18 min, return to 50% B; 18-23 min, re-equilibration at 5% B. The flow rate was 300 µL/min with an injection volume of 1µL. Mass spectrometric data were acquired on a TSQ Quantis Plus (Thermo Fisher Scientific, USA) in both positive and negative heated electrospray ionization (H-ESI) mode. Ionization parameters were as follows: spray voltage, 3.5 kV (positive) and 3.0 kV (negative); sheath gas flow, 50 a.u.; auxiliary gas flow, 10 a.u.; sweep gas flow, 1 a.u.; ion transfer tube temperature, 325°C; vaporizer temperature, 250°C. Data were acquired in profile MS mode using Xcalibur software (Thermo Fisher Scientific) and selected reaction monitoring (SRM) transitions were optimized and monitored for selected Impranil DLN-SD-derived monomers. Adipic acid was detected in negative ionization mode at m/z 145, along with its corresponding fragment at m/z 101. 1,6-hexanediol and neopentyl glycol were monitored in positive ionization mode using the m/z 119→83 and 105→87 transitions, respectively. 6-hydroxycaproate was acquired in full-scan (FS) mode (m/z 50–1000; Q1 resolution, 1.2 FWHM). Calibration curves for the quantification of adipic acid, 6-hydroxycaproate, ethylene glycol, and neopentyl glycol were constructed over a concentration range of 0.3-200 µM.

HPLC analysis was performed to quantify the amount of adipic acid, 6-hydroxycaproate, and neopentyl glycol. Prior to analysis, the supernatants were centrifuged through 0.45 µm nylon microcentrifuge filters at 13,000 rpm for 10 min at room temperature to remove any residual polymer precipitates and, when necessary, diluted prior to analysis. The HPLC system used was a Nexera Lita (Shimadzu), equipped with a Resex ROA-Organic Acid H^+^ (8 %) column (300 mm x 7.8 mm, Phenomenex). A sample volume of 5 μL was injected into the column. The mobile phase consisted of 2.5 mM H_2_SO_4_, delivered at a flow of 0.6 mL/min, with the column maintained at 60 °C. Separated components were detected by a variable wavelength detector set at 210 nm (Photodiode Array Mod. Shimadzu SPD-M40), and a refractive index detector (Shimadzu RID-20A). Peaks were identified by comparison with reference standards dissolved in ultrapure water. Calibration curves for peak quantification were constructed over a concentration range of 0.5-100 mM.

### 2.5 Protein expression and purification

Strains YM4B615 and YM4B5105, expressing *Fo*Cut5a and the *Fo*Cut5a-*Tr*CBM1 enzymes, respectively, were cultivated in 500 mL of YPD medium at 28 °C and 200 rpm for approximately 24 h. Prior to purification, supernatants were filtered through 0.22 μm polyethersulfone (PES) membrane filters (Membrane Solutions, USA), 10-fold concentrated using a 10 kDa cut-off PES ultrafiltration membrane (Merck Millipore, U.S.) and dialyzed for 16 h at 4 °C against a 50 mM Tris-HCl pH 8.0 buffer containing 300 mM NaCl. The two enzymes were purified as previously described (Makryniotis et al., 2024) while homogeneity and purity of the isolated enzymes were verified by sodium dodecyl sulphate-polyacrylamide gel electrophoresis (SDS-PAGE), using a 12.5 % polyacrylamide gel. Protein quantification was performed by measuring the absorbance at 280 nm, using the molar extinction coefficients computed by the ProtParam tool from ExPASy (https://web.expasy.org/protparam/). The fractions containing the purified enzymes were dialyzed against a 20 mM Tris-HCl pH 8.0 buffer.

### 2.6 Enzymatic activity

Transformants carrying the four esterase constructs were screened for extracellular esterolytic activity as a proxy for functional enzyme secretion. For each construct, 11-12 independent colonies were cultivated in 1 mL YPD medium in 2 mL microcentrifuge tubes for 24 h at 28 °C with shaking at 200 rpm. Culture supernatants were recovered by centrifugation and assayed using *p*-nitrophenyl butyrate (*p*NPB) as substrate. Each 250 µL reaction contained 50 mM phosphate-citrate buffer (pH 6.0), 1.1 mM *p*NPB and 20 µL of culture supernatant. The substrate was supplied from a 20 mM stock solution in DMSO (Thermo Scientific Chemicals, USA), resulting in a final DMSO concentration of 5.5% (v/v). Control reactions containing buffer, substrate and DMSO but no supernatant were included, and the corresponding absorbance change was subtracted to correct for spontaneous substrate hydrolysis. Reactions were incubated at 37 °C for 10 min, and *p*-nitrophenol (*p*NP) release was monitored at 410 nm using a Cytation™ 3 multimode microplate reader (BioTek Instruments, USA). Extracellular esterolytic activity was expressed as the amount of *p*NP released per minute and per millilitre of culture supernatant and was calculated as:

activity [mU/mL] = (ΔA_410_/Δt) × a × 10^3^ × (V_reaction_/V_supernatant_)

where ΔA_410_/Δt is the blank-corrected rate of absorbance increase (AU/min), *a* = 0.4007 mM/AU is the slope of a *p*NP calibration curve generated in assay buffer (pH 6.0) and measured under the same assay conditions, thereby accounting for both well path length and the protonation state of *p*NP, Vreaction = 250 µL, and Vsupernatant = 20 µL. The factor 10³ converts mmol/L to nmol/mL. Activities are reported as mU/mL, where 1 mU corresponds to the release of 1 nmol of *p*NP per minute. Each transformant value represents the mean of two technical replicates.

The esterolytic activity of purified *Fo*Cut5a and *Fo*Cut5a-*Tr*CBM1 was evaluated on *p*NPB, at 37 °C for 10 min. Each reaction mixture comprised of 0.05 M phosphate-citrate buffer (pH 6.0) containing 1.1 mM *p*NPB and 3.2 µg/mL of purified enzyme. The release of *p*NP was measured at 410 nm using a SpectraMax-250 microplate reader (Molecular Devices, USA) connected to SoftMaxPro software (version 1.1, Molecular Devices, US). Enzymatic activity was expressed in Units (U), defined as the amount of enzyme required to release 1 μmol of *p*NP per minute. A calibration curve was constructed for the quantification of *p*NP in the utilized buffer system, in a range of 0-167 µg/mL.

The polyurethane-degrading activity of purified *Fo*Cut5a and *Fo*Cut5a-*Tr*CBM1 was evaluated using an Impranil DLN-SD turbidity-reduction assay, as previously described by Soto-Hernández et al., (2025) . Reactions were prepared in triplicate by adding purified *Fo*Cut5a or *Fo*Cut5a-*Tr*CBM1 to a final concentration of 2 µg/mL in 1% (v/v) Impranil DLN-SD suspended in 0.1 M sodium phosphate buffer pH 7.0. Control reactions containing Impranil DLN-SD without enzyme were included to account for non-enzymatic turbidity changes. Samples were incubated at 30 °C with shaking at 1,200 rpm, and polyurethane hydrolysis was monitored by measuring the decrease in optical density at 600 nm (OD₆₀₀) using a SpectraMax-250 microplate reader (Molecular Devices, USA). The enzymatic activity was expressed as ΔOD₆₀₀/(min x mg).

Three independent enzyme purifications (biological replicates) were used for all experiments. Esterase activity was measured in technical triplicate using the *p*NPB assay, whereas Impranil DLN-SD degradation was assessed in technical triplicate by monitoring turbidity reduction.

### 2.7 Biochemical characterization

Optimum temperature was determined by assaying enzyme activity on *p*NPB, following the standard *p*NP assay conditions, over a range of temperatures (30-45 °C), while thermostability was investigated by assessing enzymes’ residual activity on *p*NPB, after their incubation in 20 mM Tris-HCl pH 8.0 buffer, at 30° C and 40° C for a duration of up to 2 h. Moreover, optimum pH was measured by defining enzyme activity on *p*NPB, following the standard *p*NP assay conditions, over a range of pH between (6 and 9). Buffer systems used were 50 mM citrate-phosphate (C-P, pH 6.0), 50 mM sodium-phosphate (S-P, pH 7.0) and 50 mM Tris-HCl (T-H, pH 8.0–9.0).

### 2.8 Data analysis

Data are presented as mean ± standard error of the mean (SEM), with the replicate structure specified in each figure legend. Biological replicates represent independent cultures or enzyme preparations, whereas technical replicates correspond to repeated measurements of the same sample and were averaged prior to statistical analysis. All tests were two-sided, with significance set at *P* < 0.05 and Holm correction applied within each family of comparisons. Statistical analyses were performed in Python 3.11.16 using SciPy 1.17.1 and statsmodels 0.15.0. Exact *P* values and the statistical test used for each analysis are indicated in the corresponding figure or figure legend. Figures were generated using Python 3.11 with Matplotlib 3.11 and NumPy 2.4.

## 3. Results

### 3.1 Identification of a functionally expressed esterase in *Yarrowia lipolytica*

To select an esterase for subsequent poly(ester-urethane)-degradation studies, four candidate enzymes were initially screened and compared: *Fo*Cut5a, *Da*PUase, *Se*1JFR, and *Apa*rEST. *Fo*Cut5a was described to act on both aliphatic and aromatic polyesters, including poly(ε-caprolactone) (PCL) and PET-related model substrates (Dimarogona et al., 2015; Nikolaivits et al., 2016). *Da*PUase is a polyurethane-degrading esterase from the bacterium *Comamonas acidovorans* that targets ester bonds within poly(ester urethanes) and related synthetic polymers (Nomura et al., 1998; Pantelic et al., 2023). *Se*1JFR is a bacterial cutinase-like enzyme from *Streptomyces cavourensis*, recently described as a PETase-like lipase with broad substrate specificity toward various aliphatic and aromatic polyesters (Makryniotis et al., 2024). *Apar*Est is a putative carboxylesterase from the filamentous fungus *Aspergillus parasiticus*, identified and selected after secretome proteomic analysis following cultivation in presence of PLA oligomers (unpublished data). Collectively, the four enzymes represent phylogenetically distinct polyesterase clades, enabling the screening of a broad range of polyester-degrading enzymes rather than closely related variants of the same enzyme family.

Each esterase was cloned between the TEF1 promoter and the LIP2 terminator and, in the same assembly reaction, fused to four different signal peptides to promote enzyme secretion, generating for each esterase a mini-library of four plasmid variants (Figure 1a). Four signal peptides previously validated (Celińska et al., 2018) for heterologous protein expression and secretion in *Y. lipolytica* were tested: SigPep1, derived from the *YALI0E22374g* locus; SigPep2, corresponding to the native lipase Lip2 signal peptide; SigPep3, derived from the native SoAMY signal peptide; and SigPep4, originating from the alkaline extracellular protease XPR2 (Table 1). The four esterase libraries were assembled by GGA, propagated in *E. coli*, and sequencing of three randomly selected clones from each esterase library confirmed correct assembly, maintenance of the reading frame, and the presence of the expected diversity. The resulting libraries were stably integrated into the F11 site of the genome of *Y. lipolytica* (Yuzbashev et al., 2023). Transformants were plated on minimal medium containing 1% glucose, and 11-12 colonies were randomly selected for further analysis. Esterase activity was quantified in the supernatants of overnight cultures using the *p*NPB assay (Figure 1b). Esterase activities of *Fo*Cut5a transformants showed a broader distribution than those of the other enzymes. While differences in median activity were limited, several individual *Fo*Cut5a clones displayed substantially higher activity than the population baseline (Figure 1b). Since this assay relies on a soluble model substrate, the observed differences primarily reflect the ability of each enzyme to be efficiently expressed, correctly secreted, and to retain catalytic activity in the *Y. lipolytica* host, rather than directly predicting performance on polymeric substrates. Based on the outcome of this initial screening, *Fo*Cut5a was selected for subsequent analyses.

**Figure 1.**
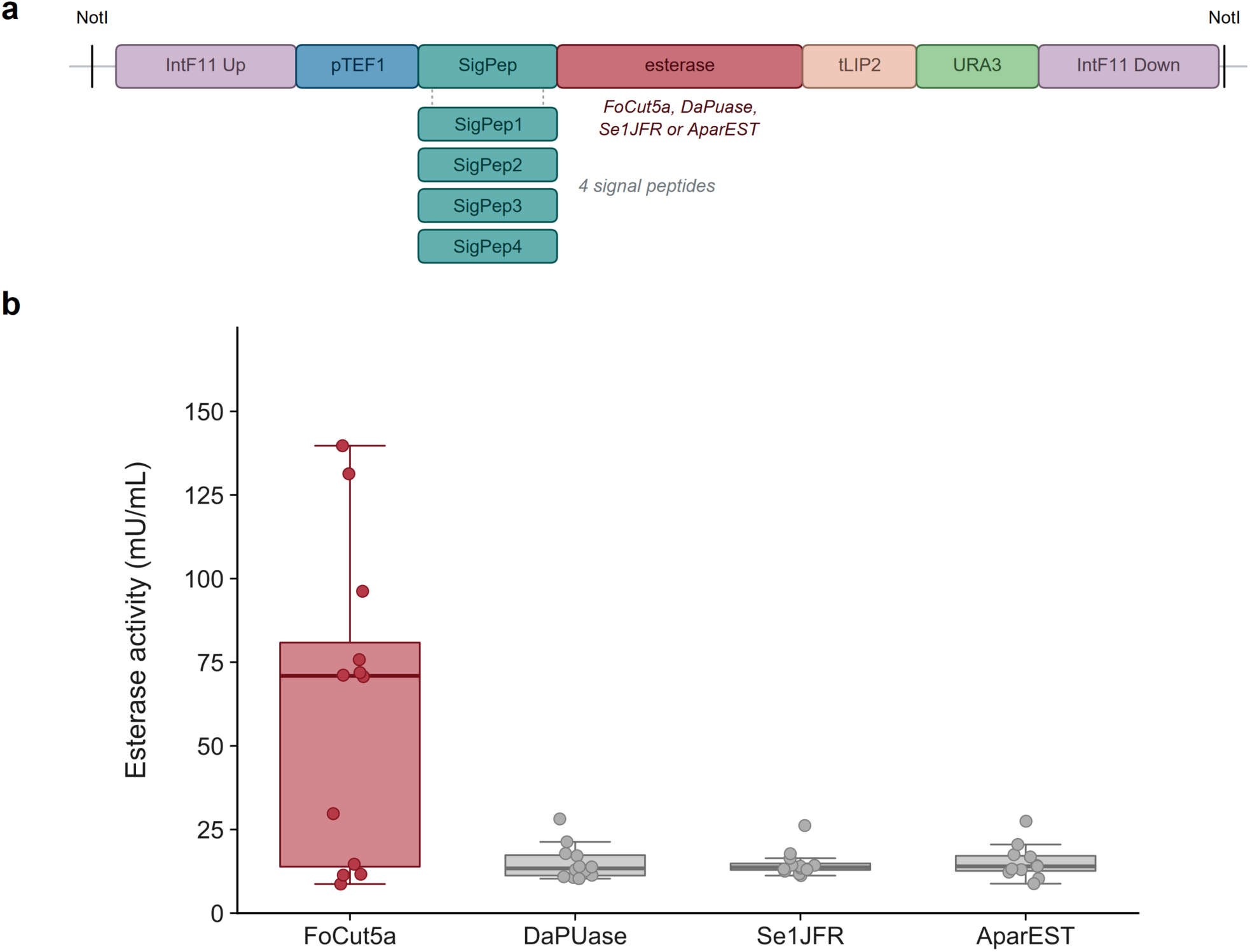
Selection of a secreted esterase in *Y. lipolytica*. **(a)** Schematic representation of the integration cassette containing promoter (pTEF1), a signal peptide (SigPep), the esterase coding sequence, terminator (tLIP2), and auxotrophic marker (URA3). The expression cassette is flanked by homology arms for site-specific integration into the F11 locus (IntF11 Up and IntF11 Rev) and by NotI restriction sites, which are used to linearize the cassette prior to transformation. Four esterases (*Fo*Cut5a, *Da*PUase, Se1JFR, and *Apar*EST) were each combined with four candidate signal peptides (SigPep1-SigPep4; teal), generating four secretion variants per esterase. **(b)** Esterase activity in culture supernatants of individual transformants, quantified by *p*-nitrophenyl butyrate (*p*NPB) hydrolysis and expressed as mU/mL, where 1 mU corresponds to the release of 1 nmol of *p*-nitrophenol (*p*NP) per minute. Each dot represents an independent transformant and corresponds to the mean of two technical replicates (n = 12 transformants for *Fo*Cut5a, *Da*PUase and Se1JFR; n = 11 for *Apar*EST). Boxes indicate the interquartile range (IQR) with the median shown as a horizontal line; whiskers extend to 1.5 × IQR, and dots represent individual transformants. Statistical significance was assessed by two-sided Mann-Whitney U tests comparing *Fo*Cut5a with each of the other esterases, followed by Holm correction for multiple testing (*Fo*Cut5a vs *Da*PUase: adjusted P = 0.068; vs Se1JFR: adjusted P = 0.091; vs *Apar*EST: adjusted P = 0.091).

### 3.2 Construction of Modular Enzyme Library for Poly(ester-urethane) Degradation

Having identified *Fo*Cut5a as the most promising candidate for expression in *Y. lipolytica*, we next employed a combinatorial synthetic biology approach to optimize its secretion and polymer-binding properties through modular protein engineering. In addition to the signal peptide variants described above, the effects of selected linker sequences and SBDs on polyurethane-degrading activity were also evaluated (Table 1). To modulate the spatial arrangement and substrate accessibility of the enzyme, four linker sequences exhibiting different structural and physicochemical properties were selected. These included Linker 1, a flexible, protease-resistant cellobiohydrolase-derived linker from *Trichoderma reesei* rich in proline, glycine, and threonine residues (Graham et al., 2022); Linker 2, the intrinsically disordered interdomain linker of the Lytic Polysaccharide Monooxygenase (LPMO) *Sc*AA10 from *Streptomyces coelicolor* (Courtade et al., 2018); Linker 3, the native serine/threonine-rich linker of the LPMO *Bc*AA9C from *Botrytis cinerea* (Srivastava et al., 2022); and Linker 4, a structured fibronectin type III-like linker derived from the polyhydroxybutyrate (PHB) depolymerase of *Delftia acidovorans* (Kasuya et al., 1997). In addition, six SBDs and surface-interacting proteins were evaluated for their potential to enhance enzyme adsorption to insoluble polyurethane substrates. These included *Tr*CBM1, a cellulose-binding module from *Trichoderma reesei* (Ribitsch et al., 2013); *Tt*CBM10, a type A carbohydrate-binding module from *Teredinibacter turnerae* (Graham et al., 2022); PBM, a polyhydroxyalkanoate-binding module from *Alcaligenes faecalis* (Ribitsch et al., 2013); PHB, the substrate-binding domain of the PHB depolymerase from *Delftia acidovorans* (Shinomiya et al., 1998); and the class II Hydrophobins 4 (HFB4) and 7 (HFB7) from *Trichoderma reesei* and *Trichoderma harzianum*, respectively (Ribitsch et al., 2015). All SBDs fusion constructs were engineered with a C-terminal V5-His₆ tag to facilitate affinity purification.

To explore how modular architecture influences poly(ester-urethane)-degrading performance, a 96-construct combinatorial library was assembled and overexpressed in *Yarrowia lipolytica* using CRISPR/Cas9-mediated genome editing via the *YaliCraft®* system (Yuzbashev et al., 2023). The library was constructed via BsaI-mediated GGA by systematically varying four classes of functional elements. Each construct integrated one of four signal peptides, a single catalytic domain-cutinase *Fo*Cut5a from *Fusarium oxysporum*, one of four linker modules, and one of six SBDs, generating all 96 possible combinations. The library incorporated the integration backbone together with the TEF1 promoter and LIP2 terminator (Figure 2). The library was propagated in *E. coli*, and sequencing of six randomly selected clones confirmed correct assembly, preservation of the reading frame, and representation of the expected combinatorial diversity. The library of expression cassettes was released by NotI digestion and subsequently integrated into the F11 locus of *Y. lipolytica* Y-4973 using a CRISPR/Cas9-mediated strategy (Yuzbashev et al., 2023). Transformants were selected on minimal medium containing 1% Impranil DLN-SD as the sole carbon source, allowing direct screening for poly(ester-urethane)-degrading activity. Four independent transformation experiments yielded 142 colonies on Impranil plates, corresponding to ∼1.5-fold coverage of the 96-member library (148%).

**Figure 2.**
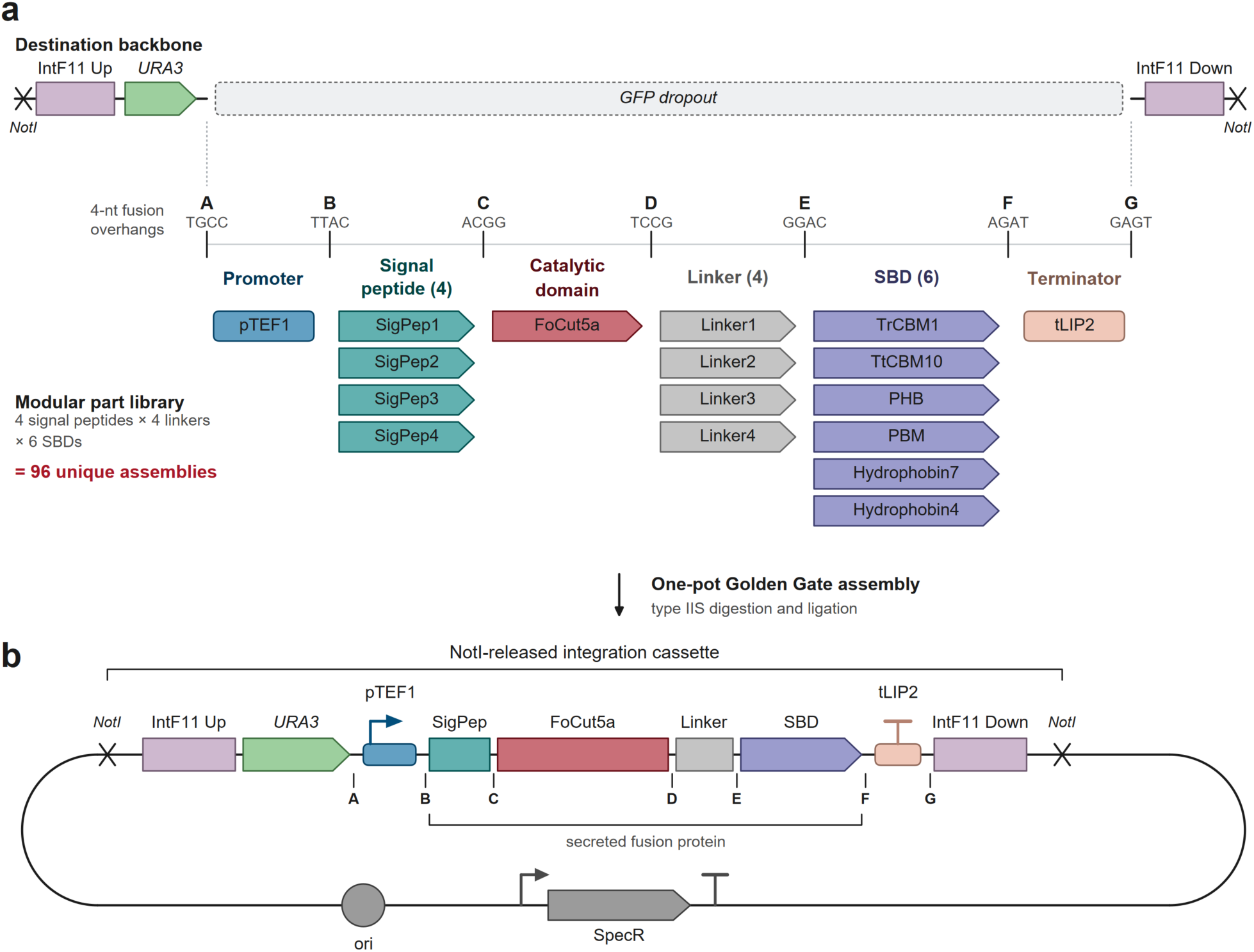
**(a)** Design of the combinatorial Golden Gate library. A destination backbone containing a green fluorescent protein (GFP) dropout cassette was combined in a one-pot Golden Gate assembly with a collection of modular parts comprising one promoter (pTEF1), four signal peptides (SigPep), one catalytic module (*Fo*Cut5a), four linkers, six substrate-binding domains (SBDs), and one terminator (tLIP2). During assembly, the GFP dropout cassette is replaced by one module from each category, generating 96 possible expression constructs. The modules are flanked by defined 4-nt fusion overhangs (A-G) that direct their ordered assembly into the final expression cassette. The destination backbone additionally contains the auxotrophic marker *URA3*, homology arms for site-specific integration into the F11 locus (IntF11 Up and IntF11 Down), and NotI restriction sites used to linearize the integration cassette prior to transformation. **(b)** Architecture of the assembled integration cassette. The NotI-excised fragment contains the *URA3* marker, the expression cassette, and the IntF11 homology arms required for genomic integration. The fusion protein coding sequence joins the signal peptide, *Fo*Cut5a, linker, and SBD in frame, whereas the replication origin (ori) and spectinomycin-resistance marker (SpecR) remain on the plasmid backbone.

### 3.3 Analysis of *Fo*Cut5a Construct Combinations Enriched During Combinatorial Library Screening

To identify enriched SigPep-*Fo*Cut5a-Linker-SBD variants, all 142 transformants were screened on Impranil DLN-SD plates, and 49 colonies displaying the largest hydrolysis halos were selected for further analysis (Figure 3a). The integrated expression cassettes were amplified from genomic DNA and sequenced to determine the assembled construct. Module occurrence frequencies were subsequently calculated to evaluate the representation of individual signal peptides, linkers, and SBDs among the best-performing variants (Figure 3b-d).

**Figure 3.**
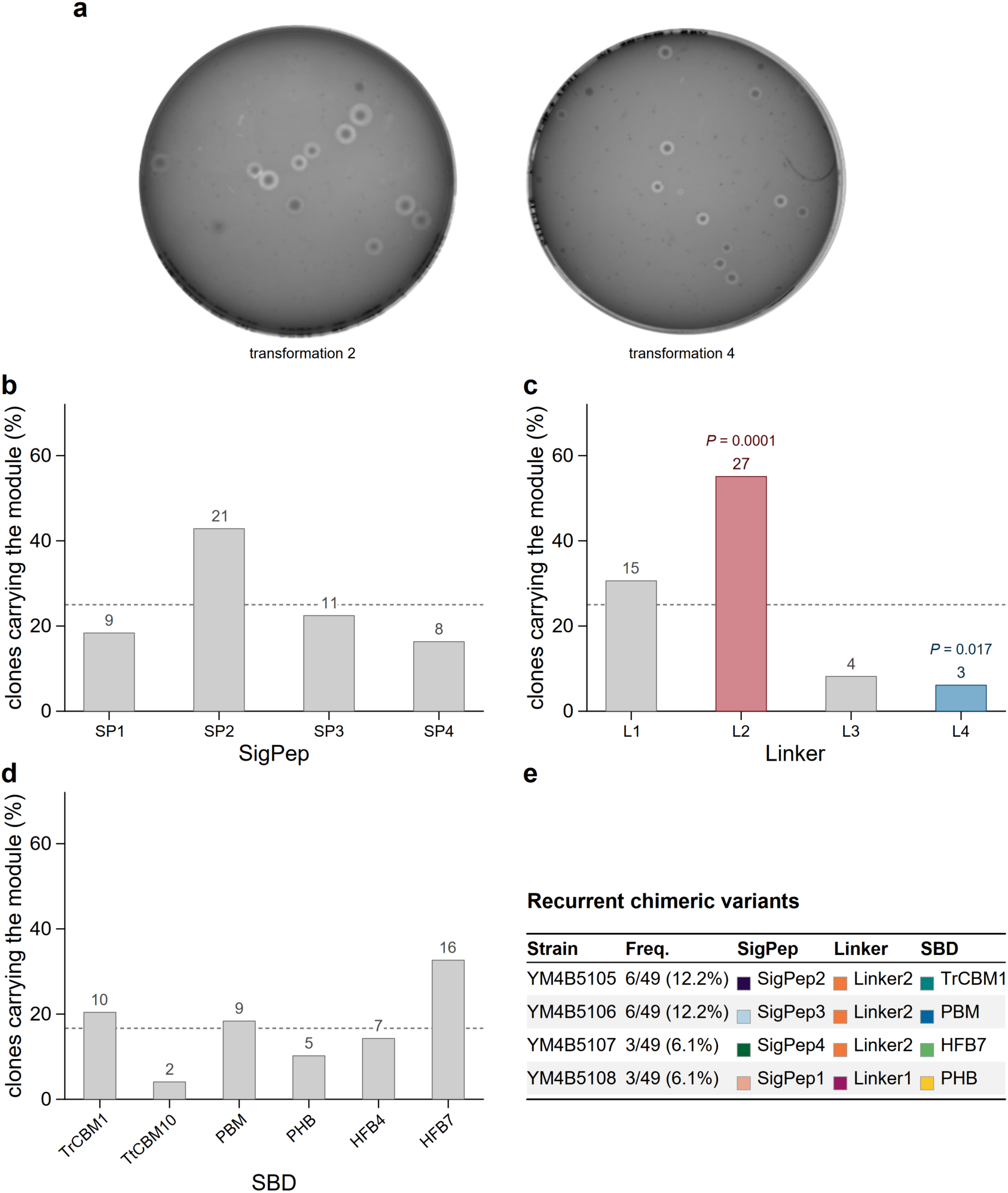
**(a)** Primary screening on 1% Impranil DLN-SD agar; representative plates from transformations 2 and 4. Frequencies of signal peptides (SigPep) **(b)**, Linkers **(c)**, and SBDs **(d)** among the 49 genotyped variants. Dashed lines indicate the frequencies expected under random sampling of the library (25% for SigPep and Linkers, 16.7% for SBDs). Module frequencies were compared with the expected distribution using two-sided exact binomial tests (n=49), followed by Holm correction for multiple testing across the 14 modules analyzed. Only two modules showed significant deviations from expectation: Linker 2 was significantly enriched (27/49 clones, 55.1%; Holm-adjusted P = 1.0 × 10⁻⁴), whereas Linker 4 was significantly underrepresented (3/49 clones, 6.1%; Holm-adjusted P = 0.017). **(e)** Most frequent chimeric variants recovered among the selected clones. The 49 genotyped clones represented 22 of the 96 possible module combinations.

Among the four signal peptides analyzed, the native *Y. lipolytica* Lip2 signal peptide (SigPep2) was the most frequently identified module, occurring in 21 of the 49 selected clones (Figure 3b). Linker 2 was the predominant linker variant, being present in 27 clones and representing the only significantly enriched module relative to its expected frequency in the library (P = 0.0001) (Figure 3c). Among the SBDs, HFB7 was the most prevalent (16 occurrences), followed by *Tr*CBM1 and PBM, which were identified in 10 and 9 clones, respectively (Figure 3d).

The strong overrepresentation of Linker 2 suggests that flexible and intrinsically disordered interdomain connectors may be advantageous for *Fo*Cut5a activity on Impranil DLN-SD. Linker 2 (30 aa) and Linker 1 (24 aa) are both enriched in threonine, proline, and glycine residues, which together represent 70% and 71% of their amino acid compositions, respectively, and have been reported to function as extended molecular tethers that facilitate the independent movement of catalytic and binding domains (Courtade et al., 2018; Graham et al., 2022). Such architectures could facilitate exploration of the poly(ester-urethane) surface by the catalytic domain while maintaining substrate association through the binding module.

The most frequently recovered SBDs were HFB7, *Tr*CBM1, and PBM (Figure 3d) and have previously been associated with enhanced polyester or polyurethane degradation. HFB7 mediates amphiphilic surface interactions that may facilitate enzyme adsorption to hydrophobic polymer substrates (Espino-Rammer et al., 2013; Ribitsch et al., 2015), *Tr*CBM1 promotes interactions with hydrophobic polyester surfaces (Ribitsch et al., 2013), and PBM has been shown to improve substrate association and hydrolytic performance in polyurethane-active enzymes (Gamerith et al., 2016). Their frequent occurrence among the highest-performing clones is therefore consistent with a beneficial role of substrate targeting during Impranil DLN-SD degradation.

Analysis of complete module combinations provided additional insight beyond the frequencies of individual elements. Among the 96 theoretical SigPep-*Fo*Cut5a-Linker-SBD combinations present in the library, only 22 were recovered among the 49 selected clones (Figure 3e). Random sampling of 49 clones from the library would be expected to yield approximately 38.5 distinct combinations, compared with 22 combinations recovered among the selected clones. Several combinations were repeatedly identified, including SigPep2-Linker2-*Tr*CBM1 (6/49 clones), SigPep3-Linker2-PBM (6/49), SigPep4-Linker2-HFB7 (3/49), and SigPep1-Linker1-PHB (3/49), suggesting that specific module assemblages may confer a selective advantage under the screening conditions employed. Representative strains carrying recurrent combinations (Figure 3e) were subsequently selected for re-sequencing and further characterization.

### 3.4 Monomer Release and Metabolic Conversion during Impranil DLN-SD Degradation by *Y. lipolytica*

Despite the proprietary nature of Impranil DLN-SD, its chemical architecture has been inferred from degradation studies. Biffinger et al., (2015) proposed that the material consists of polyester domains formed from adipic acid, 1,6-hexanediol, and neopentyl glycol, linked through urethane bonds derived from hexamethylene diisocyanate (HDI). Subsequent microbial and enzymatic degradation experiments have independently corroborated this model through the detection of the corresponding monomeric constituents released during polymer breakdown (Fuentes-Jaime et al., 2022; de Witt et al., 2024). In addition, Fuentes-Jaime et al., (2022) detected HDI, further supporting the presence of urethane segments derived from this diisocyanate in the poly(ester-urethane) structure.

As the release of these degradation products has not yet been investigated in a yeast-based system, they were selected as marker compounds to assess PU degradation by *Y. lipolytica*. To this end, strain YM4B5105 (Figure 3e) was selected for monomer analysis from the four constructs producing the largest hydrolysis halos, as it represented one of the most frequently recovered architectures and combined several of the features that emerged as enriched during screening, namely SigPep2, Linker2, and *Tr*CBM1 (Figure 3).

Cells were transferred from overnight YPD cultures to minimal medium containing 1% Impranil DLN-SD at an initial OD_600_ of 1.0 and incubated at 28 °C with shaking at 200 rpm for 144 h. Samples were collected at selected time points during incubation, and culture supernatants were subsequently analyzed by LC-MS to identify and quantify monomers released from polyurethane degradation. Biomass remained essentially unchanged throughout the incubation period, with no significant increase relative to the initial biomass. In parallel, an abiotic control consisting of Impranil DLN-SD-containing minimal medium without added cells was included. No detectable degradation products were observed in this control by LC-MS analysis (data not shown). In contrast, cultures of the *Y. lipolytica* strain expressing *Fo*Cut5a produced adipic acid, neopentyl glycol, and 1,6-hexanediol as the major degradation products (Figure S1a–c). Adipic acid and neopentyl glycol accumulated progressively throughout the incubation, reaching 1.96 ± 0.21 mM and 3.43 ± 0.53 mM, respectively, after 144 h (Figure 4a). By contrast, 1,6-hexanediol exhibited a transient accumulation pattern, reaching a maximum concentration of 1.16 ± 0.32 mM after 7 h, followed by a progressive decline to near-background levels by 144 h (Figure 4b).

**Figure 4.**
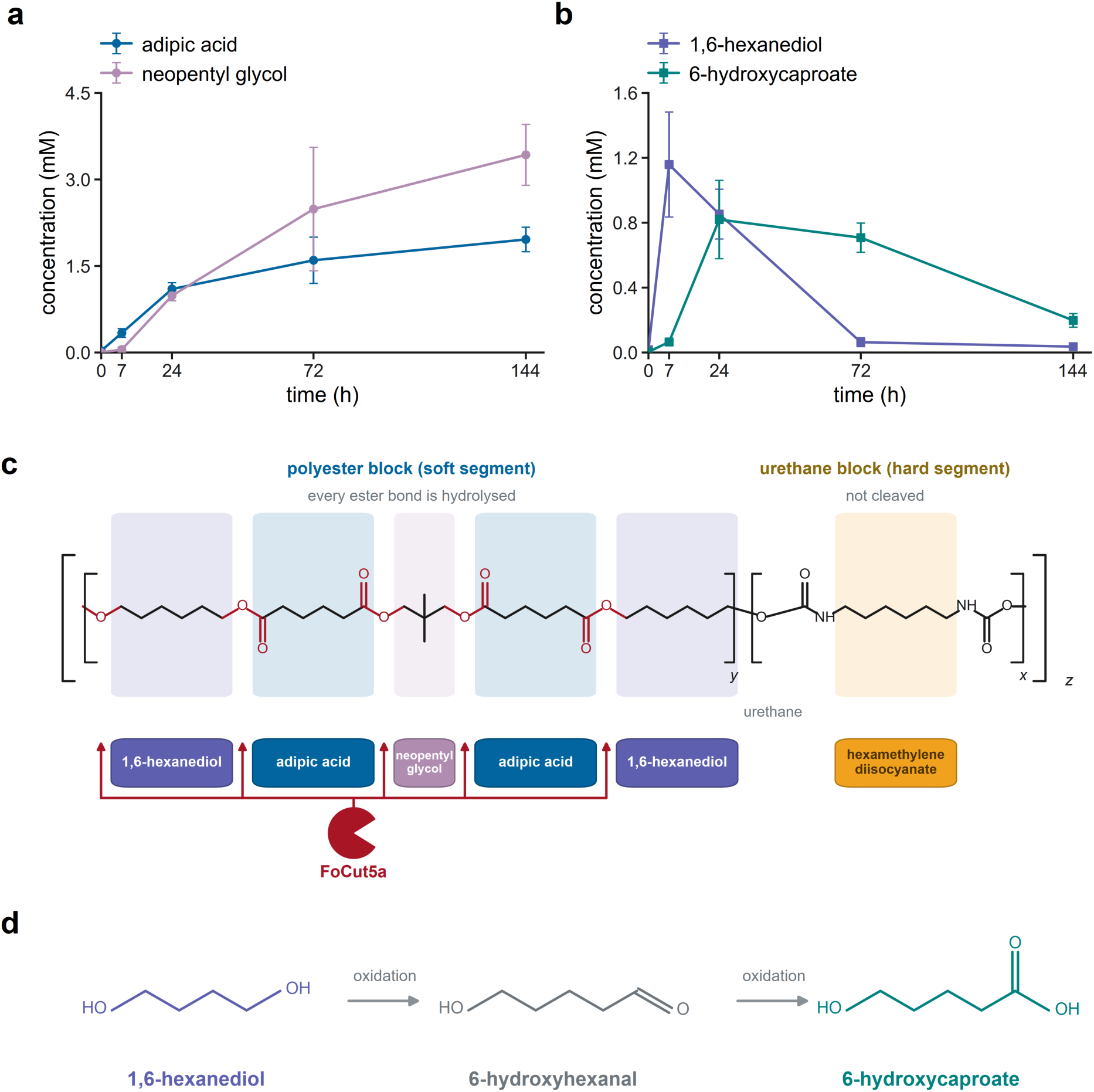
**(a,b)** Release of poly(ester-urethane)-derived monomers from Impranil DLN-SD by *Y. lipolytica* strain YM4B5105 expressing *Fo*Cut5a. **(a)** Accumulation of adipic acid (blue) and neopentyl glycol (mauve), and **(b)** 1,6-hexanediol (violet) and 6-hydroxycaproate (teal), quantified by LC-MS after 0, 7, 24, 72, and 144 h of cultivation on 1% Impranil DLN-SD. Data points represent the mean of 2-5 independent measurements (n varied depending on the analyte and time point), and error bars indicate the standard error of the mean (SEM). **(c)** Proposed structure of Impranil DLN-SD, consisting of alternating polyester soft segments (y) and urethane hard segments (x) repeated along the poly(ester-urethane) chain (z). Monomeric units are colour-coded as in panels a and b; hexamethylene diisocyanate (amber) was not detected among the degradation products. Ester bonds (red) indicate the *Fo*Cut5a cleavage sites (arrows), releasing adipic acid, 1,6-hexanediol, and neopentyl glycol. The diol connecting soft and hard segments remains linked to the hard segment because it is flanked by an ester bond and a urethane bond. No carbamate bond hydrolysis was observed, consistent with the absence of hexamethylene diisocyanate in the culture supernatant. **(d)** Proposed metabolic conversion of 1,6-hexanediol to 6-hydroxycaproate via the intermediate 6-hydroxyhexanal.

This product profile is consistent with selective degradation of the polyester soft segment of Impranil DLN-SD, as cutinases hydrolyze ester bonds but not urethane (carbamate) linkages, whose cleavage requires amidase activity (Schmidt et al., 2017; Magnin et al., 2019). Accordingly, degradation is expected to be largely confined to the polyester domain, leaving the hard segment unaffected. The release of adipic acid, neopentyl glycol, and 1,6-hexanediol matches the proposed composition of the soft segment and provides direct evidence of extensive ester-bond hydrolysis within this region. Consistent with this mechanism, the diol linking the soft and hard segments is expected to remain attached to the hard segment because it is bounded by an ester bond on one side and a urethane bond on the other (Figure 4c). Likewise, no evidence of carbamate-bond hydrolysis was obtained, as no HDI-derived compounds were detected in the culture supernatant, unlike the findings of Fuentes-Jaime et al., (2022).

Moreover, given the rapid depletion of 1,6-hexanediol (Figure 4b), we investigated whether its consumption was associated with the formation of downstream metabolites in the culture medium. We specifically searched for 6-hydroxycaproate, which can arise from the oxidation of one terminal hydroxyl group of 1,6-hexanediol. Previous studies have shown that *Y. lipolytica* converts ethylene glycol (Messina et al., 2026) and 1,4-butanediol (Carniel et al., 2023) into their corresponding hydroxy acids. In contrast, Carniel et al., (2023) reported extensive consumption of 1,6-hexanediol without detecting corresponding oxidation products under the analytical conditions employed. Our LC-MS analysis revealed the presence of 6-hydroxycaproate, a compound not previously reported among the products derived from Impranil DLN-SD (Figure S1d). Its concentration increased rapidly, becoming detectable after 7 h and reaching 0.82 ± 0.24 mM at 24 h, before declining to 0.20 ± 0.04 mM at 144 h (Figure 4b). Together with the earlier appearance of 1,6-hexanediol, this pattern is consistent with the conversion of the polymer-derived diol into 6-hydroxycaproate via the intermediate 6-hydroxyhexanal (Figure 4d), likely catalyzed by a fatty alcohol oxidase and a fatty aldehyde dehydrogenase, as previously proposed for aliphatic alcohol metabolism in *Y. lipolytica* (Carniel et al., 2023). The transient accumulation of 6-hydroxycaproate, followed by its decline, suggests further metabolism rather than assimilation for growth, as biomass remained unchanged over the same period. Although conversion to adipic acid cannot be excluded, the metabolite profiles provide little support for this hypothesis, since adipic acid concentrations remained largely stable between 72 and 144 h, when the sharpest decrease in 6-hydroxycaproate was observed (Figure 4a, b).

To our knowledge, these findings provide the first experimental evidence that *Y. lipolytica* converts 1,6-hexanediol into 6-hydroxycaproate during poly(ester-urethane) degradation. These results reveal a previously unrecognized metabolic route linking the release of polyester-derived monomers to microbial metabolism through the oxidation of polymer-derived 1,6-hexanediol.

### 3.5 Screening of Engineered *Y. lipolytica* Strains for Enhanced Polyurethane Degradation

To assess whether specific combinations of SigPep, linker, and SBD influenced the cleavage of ester bonds within the polyester domains of Impranil DLN-SD, the four strains carrying different *Fo*Cut5a-derived constructs (Figure 3e) were cultivated in minimal medium containing 1% Impranil DLN-SD at an initial OD₆₀₀ of 1. Culture supernatants were collected at multiple time points over the 144 h incubation period and analyzed by HPLC to quantify the major products generated during polyurethane degradation. In addition to the four engineered strains expressing different *Fo*Cut5a variants, the control strain YM4B5102, carrying the empty destination backbone used in the GGA platform (Figure 2a), was also included in the analysis.

HPLC analysis revealed the accumulation of adipic acid and 6-hydroxycaproate in all cultures supplemented with Impranil DLN-SD (Figure 5), while biomass remained essentially unchanged throughout the experiment. Notably, the control strain YM4B5102 also released both compounds, accumulating 2.2 ± 0.02 mM adipic acid after 144 h, whereas 6-hydroxycaproate, which is formed by oxidation of the released 1,6-hexanediol, reached a maximum concentration of 0.8 ± 0.09 mM after 24 h before subsequently declining (Figure 5). These findings demonstrate that *Y. lipolytica* possesses an endogenous capacity to cleave ester bonds within the polyester domains of Impranil DLN-SD, likely through the action of secreted esterases. Consistent with its saprophytic lifestyle, *Y. lipolytica* secretes Lip2 together with several extracellular hydrolytic enzymes (Pignède et al., 2000; Fickers et al., 2005). Notably, Lip2 overexpression has recently been employed as a strategy to improve polyester-degrading capabilities in engineered strains (Dybka & Mirończuk, 2026).

**Figure 5.**
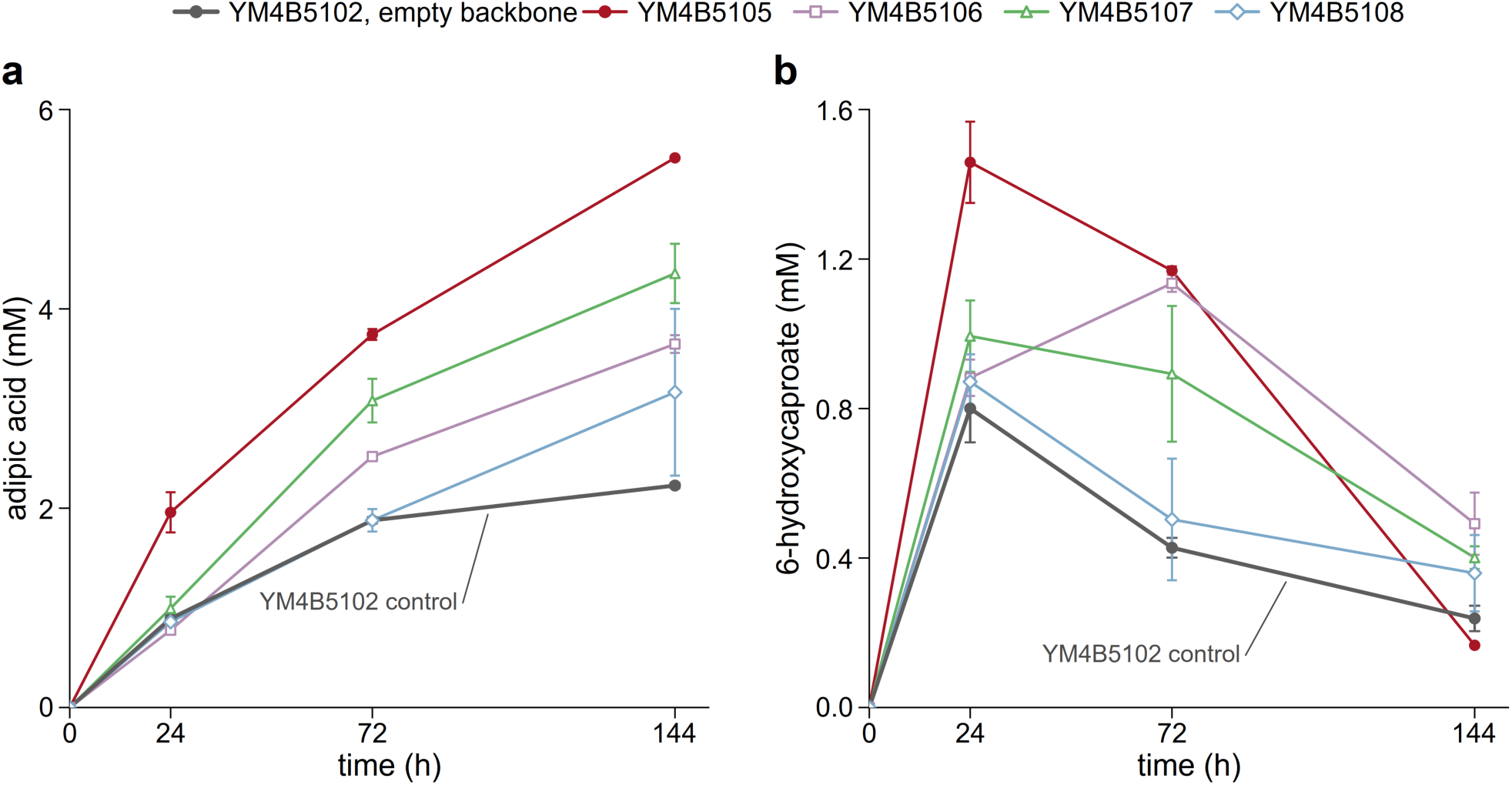
Polyurethane degradation products released by engineered and control strains. *Y. lipolytica* strains were cultivated in YNB supplemented with 1% Impranil DLN-SD, and the concentrations of adipic acid **(a)** and 6-hydroxycaproate **(b)** in culture supernatants were monitored by HPLC over 144 h. The empty-backbone control strain YM4B5102 is shown in grey, while engineered strains YM4B5105, YM4B5106, YM4B5107, and YM4B5108 are shown in dark red, purple, green, and blue, respectively. Data points represent the mean of 2-5 independent measurements (n varied depending on the analyte and time point), and error bars indicate the standard error of the mean (SEM).

Nevertheless, expression of the *Fo*Cut5a variants substantially enhanced ester-bond cleavage. Adipic acid accumulated progressively in all strains and reached 5.50 ± 0.04 mM in YM4B5105 after 144 h (Figure 5a). Likewise, 6-hydroxycaproate levels were higher in the engineered strains, peaking at 1.46 ± 0.11 mM in YM4B5105 after 24 h (Figure 5b). In all strains, 6-hydroxycaproate exhibited a transient accumulation profile, with maximum concentrations observed between 24 and 72 h, followed by a decline consistent with its further metabolism, as indicated by the LC-MS analysis (Figure 4b). In contrast to the LC-MS analysis, neither neopentyl glycol nor 1,6-hexanediol could be detected by HPLC. Their absence is likely due to the lower sensitivity of the refractive index detector (RID), which limits the detection of these aliphatic diols at the concentrations present in the culture supernatants.

Overall, YM4B5105, expressing *Fo*Cut5a-*Tr*CBM1 (Figure 3e), showed the highest accumulation of adipic acid and of 6-hydroxycaproate, the downstream oxidation product of the released 1,6-hexanediol (Figures 4 and 5). Both metabolites accumulated to levels exceeding those observed in the control strain YM4B5102 and in strains expressing alternative *Fo*Cut5a variants. These results identify *Fo*Cut5a-*Tr*CBM1 as the most effective construct for promoting ester-bond cleavage within the polyester domains of Impranil DLN-SD, and YM4B5105 was therefore selected for further characterization.

### 3.6 Effect of *Tr*CBM1 fusion on *Fo*Cut5a-mediated polyurethane degradation

Based on the whole-cell degradation assays, YM4B5105 was selected as the best-performing strain, as it consistently yielded the highest levels of adipic acid and 6-hydroxycaproate among all engineered variants (Figure 5). However, differences observed at the whole-cell level may not exclusively reflect differences in catalytic performance, as they can also arise from variations in enzyme expression, secretion efficiency, or protein folding.

YM4B5105 expresses a chimeric *Fo*Cut5a variant comprising (i) SigPep2, the native signal peptide of the *Y. lipolytica* lipase Lip2 (Celińska et al., 2018); (ii) Linker 2, an intrinsically disordered interdomain linker derived from the LPMO *Sc*AA10 of *Streptomyces coelicolor* (Courtade et al., 2018); and (iii) *Tr*CBM1, a carbohydrate-binding module previously shown to enhance enzyme adsorption onto polymeric substrates (Ribitsch et al., 2013). Notably, all three components were among the most frequently represented elements identified during the screening of the 49 selected colonies (Figure 3), suggesting that they may provide a functional advantage for *Fo*Cut5a-mediated degradation of Impranil DLN-SD.

Enzyme adsorption onto insoluble polymer surfaces has been identified as a major factor limiting the cleavage of ester bonds in polyester-containing substrates (Gamerith et al., 2016; Biundo et al., 2018). Accordingly, fusion of polymer-binding domains has been shown to improve degradation of insoluble polymers, including polyurethane-based materials and PET, by strengthening enzyme-substrate interactions (Ribitsch et al., 2013; Espino-Rammer et al., 2013; Gamerith et al., 2016). In this context, *Tr*CBM1 is of particular interest, as its incorporation may increase enzyme retention at the polymer surface and thereby enhance cleavage of ester bonds within the polyester domains of Impranil DLN-SD. To our knowledge, the effect of a CBM-fused fungal cutinase secreted by an engineered yeast on the degradation of a poly(ester-urethane) substrate has not been investigated previously. Whether the incorporation of *Tr*CBM1 contributes to enhanced degradation through improved adsorption to the poly(ester-urethane) surface remains to be established. To address this question, the *Fo*Cut5a-*Tr*CBM1 construct was further compared with *Fo*Cut5a lacking the SBD.

To this end, a control strain was constructed in which *Fo*Cut5a, preceded only by SigPep2 and lacking both the linker and the SBD, was integrated into the *Y. lipolytica* genome (strain YM4B615; pTEF1-SigPep2-*Fo*Cut5a-tLIP2). This newly constructed strain, together with YM4B5105, was cultivated in minimal medium supplemented with 1% Impranil DLN-SD, using an initial OD₆₀₀ of 1. HPLC analysis of culture supernatants revealed clear differences between the two strains after only 24 h of incubation (Figure 6a, b). Adipic acid accumulated to 1.96 ± 0.20 mM in YM4B5105, significantly higher than in YM4B615 (1.07 ± 0.07 mM) (Figure 6a). Likewise, 6-hydroxycaproate reached 1.46 ± 0.11 mM in YM4B5105, compared with 0.94 ± 0.12 mM in YM4B615 (Figure 6b). Both differences were statistically significant (*P* = 0.018).

**Figure 6.**
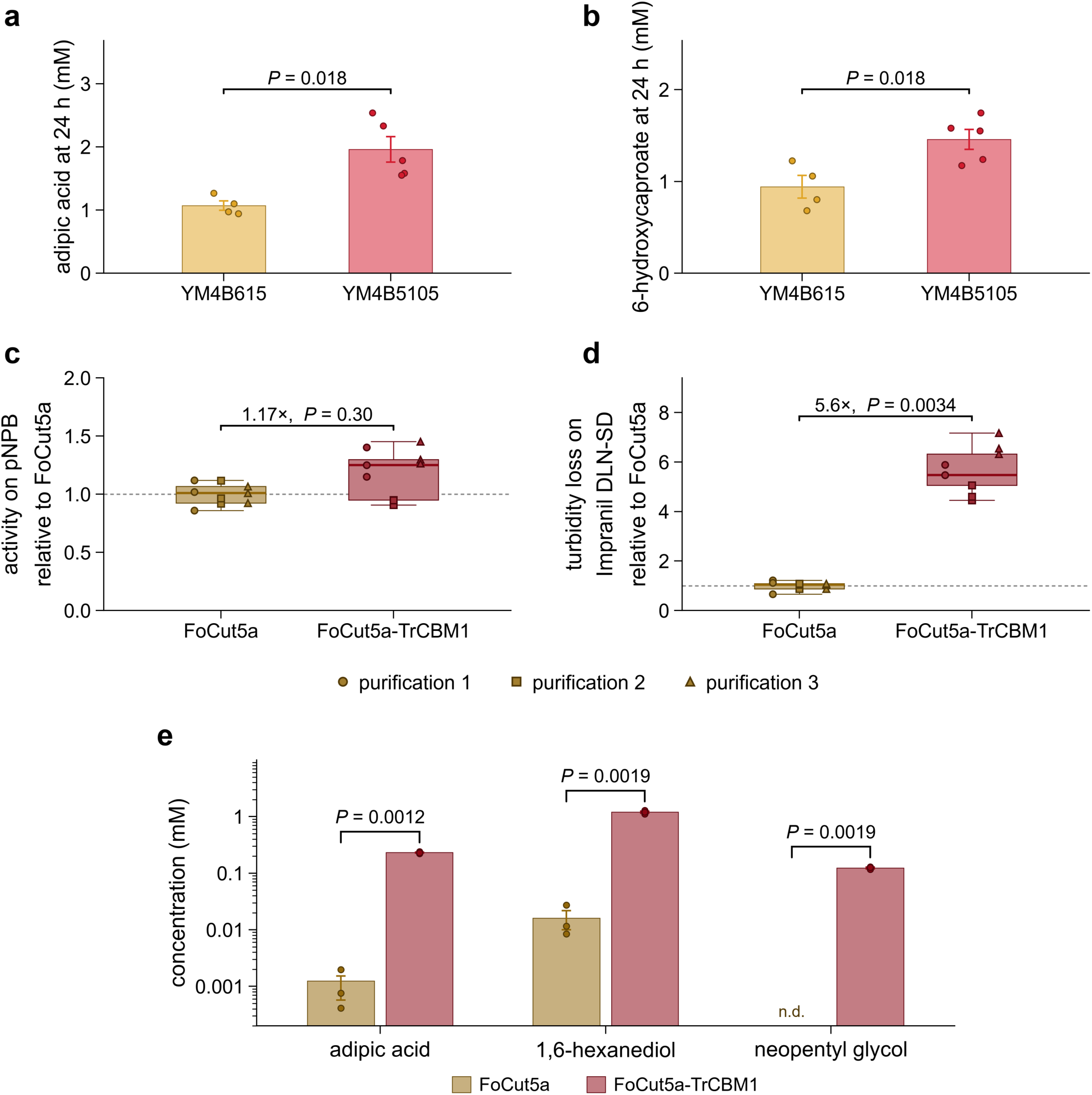
Monomer release by engineered strains and purified enzymes. **(a,b)** Adipic acid (a) and 6-hydroxycaproate (b) concentrations in culture supernatants of strain YM4B615, expressing *Fo*Cut5a, and strain YM4B5105, expressing *Fo*Cut5a-*Tr*CBM1, after 24 h of growth on 1% Impranil DLN-SD. Bars represent means, points indicate individual cultures, and error bars show the standard error of the mean (SEM) (YM4B615, *n* = 4; YM4B5105, *n* = 5). YM4B5105 accumulated significantly higher levels of both metabolites (two-sided Welch’s *t*-test with Holm correction across the two metabolites; adjusted *P* = 0.018 for both comparisons, as indicated by brackets; unadjusted *P* = 0.009 and 0.018). **(c,d)** Activity of purified *Fo*Cut5a and *Fo*Cut5a-*Tr*CBM1 on *p*-nitrophenyl butyrate (*p*NPB) (c) and Impranil DLN-SD (d). Enzymes were assayed at 3.2 µg/mL in the *p*NPB assay and 2 µg/mL in the Impranil DLN-SD turbidity assay. Activities are expressed relative to *Fo*Cut5a within each purification (dashed line = 1). Boxes indicate the median and interquartile range (IQR), whiskers extend to 1.5 × IQR, and all nine measurements per enzyme (three purifications × three technical replicates) are shown, with symbols denoting individual purifications. Statistical analysis was performed on purification means using paired two-sided *t*-tests on log₂-transformed ratios. Activities on *p*NPB did not differ significantly between the two enzymes (geometric mean ratio = 1.17, *P* = 0.30), whereas *Fo*Cut5a-*Tr*CBM1 reduced Impranil DLN-SD turbidity 5.6-fold faster (*P* = 0.0034). **(e)** LC-MS quantification of monomers released after 4 h incubation of purified enzymes with 1% Impranil DLN-SD; note the logarithmic y-axis. Bars represent means of three technical replicates, points indicate individual measurements, and error bars show SEM; n.d., not detected. Two-sided Welch’s *t*-tests with Holm correction yielded the adjusted *P* values indicated by brackets: *P* = 0.0012 for adipic acid and *P* = 0.0019 for both 1,6-hexanediol and neopentyl glycol. Gold, *Fo*Cut5a (strain YM4B615 in a,b; purified enzyme in c-e); red, *Fo*Cut5a-*Tr*CBM1 (strain YM4B5105 in a,b; purified enzyme in c-e). Darker shades in c-e denote purified enzymes.

Finally, both enzyme variants, SigPep2-*Fo*Cut5a (*Fo*Cut5a) and SigPep2-*Fo*Cut5a-Linker2-*Tr*CBM1 (*Fo*Cut5a-*Tr*CBM1), were purified to confirm their catalytic activity and to determine whether fusion to the linker and the binding module enhances cutinase activity in solution. This approach enabled the effects of biochemical activity to be evaluated independently of potential differences in expression, secretion, or protein abundance observed in vivo. Analysis of the purified enzyme preparations by 12.5% SDS-PAGE showed a single predominant band for each protein, with apparent molecular masses of approximately 24 kDa for *Fo*Cut5a and 40 kDa for *Fo*Cut5a-*Tr*CBM1 (Figure S2). While the observed molecular mass of *Fo*Cut5a was in agreement with the predicted value, *Fo*Cut5a-*Tr*CBM1 displayed an apparent molecular mass higher than the 35 kDa predicted by ExPASy ProtParam, consistent with glycosylation by the *Y. lipolytica* expression system.

The hydrolytic activity of *Fo*Cut5a and *Fo*Cut5a-*Tr*CBM1 was evaluated using a *p*NPB assay (Figure 6c) at the same protein concentration (3.2 μg/mL). Activities were measured in technical triplicate from three independent enzyme purifications. Both proteins were active in all assays, confirming correct folding and functionality. When activities were compared within each purification batch, *Fo*Cut5a-*Tr*CBM1 displayed a modest 1.17-fold increase relative to *Fo*Cut5a; however, this difference was not statistically significant (*P* = 0.30) (Figure 6c), indicating that fusion to *Tr*CBM1 does not substantially affect the intrinsic esterase activity of the enzyme toward the soluble substrate *p*NPB.

To determine whether the superior degradation phenotype of YM4B5105 (Figure 5) reflected enhanced cleavage of ester bonds within the polyester domains of Impranil DLN-SD rather than higher intrinsic esterase activity, purified *Fo*Cut5a and *Fo*Cut5a-*Tr*CBM1 were compared using a turbidity-reduction assay (Figure 6d). As enzymatic cleavage of ester bonds disrupts the light-scattering colloidal structure of Impranil DLN-SD, the resulting loss of turbidity provides a proxy for poly(ester-urethane) depolymerization and enables direct assessment of the contribution of the *Tr*CBM1 module to substrate degradation. The assay was initiated using the same enzyme concentration for both proteins (2 μg/mL), and turbidity reduction was quantified after 4 h of incubation. Under these conditions, *Fo*Cut5a-*Tr*CBM1 consistently induced a greater reduction in turbidity than *Fo*Cut5a across three independent enzyme purifications. When normalized to *Fo*Cut5a within each purification batch, the fusion enzyme exhibited a 5.6-fold higher turbidity-reduction rate (*P* = 0.0034) (Figure 6d), indicating that the *Tr*CBM1 module markedly enhances degradation of the polyester-polyurethane substrate.

To verify if the reduction in turbidity reflected enhanced cleavage of ester bonds within the polyester domains of Impranil DLN-SD, reaction supernatants collected after 4 h were analyzed by LC-MS (Figure 6e). Consistent with the turbidity assay, *Fo*Cut5a-*Tr*CBM1 released substantially higher amounts of the polyurethane-derived monomers adipic acid, 1,6-hexanediol, and neopentyl glycol than *Fo*Cut5a. *Fo*Cut5a-*Tr*CBM1 released 1.20 ± 0.05 mM 1,6-hexanediol, 0.231 ± 0.005 mM adipic acid, and 0.124 ± 0.004 mM neopentyl glycol. Under the same conditions, an equivalent amount of *Fo*Cut5a released only 16 ± 6 µM 1,6-hexanediol and 1.0 ± 0.5 µM adipic acid, corresponding to approximately 76-fold and 220-fold lower concentrations, respectively, while neopentyl glycol remained below the limit of detection (Figure 6e). Notably, 6-hydroxycaproate was not detected in reactions containing either purified enzyme, supporting the conclusion that its detection during whole-cell degradation experiments results from the oxidation of 1,6-hexanediol by *Y. lipolytica* rather than direct enzymatic release from the poly(ester-urethane).

To assess whether fusion to *Tr*CBM1 affected the biochemical properties of *Fo*Cut5a, enzyme activity was measured across different pH values and temperatures using *p*NPB as substrate. Both *Fo*Cut5a and *Fo*Cut5a-*Tr*CBM1 showed maximal activity at pH 8.0 and 40 °C, indicating that the fusion did not alter the optimal conditions (Figure S3a, b). Thermal stability assays revealed that both enzymes were highly thermolabile. At 30 °C, residual activity dropped below 14% after 90 min and was completely lost after 120 min. At 40 °C, no activity was detected after 60 min (Figure S3c). These results indicate that the improved polyurethane hydrolysis observed for *Fo*Cut5a-*Tr*CBM1 is not attributable to enhanced thermal stability.

Collectively, these results indicate that fusion of the *Tr*CBM1 binding domain does not significantly enhance the intrinsic catalytic activity of *Fo*Cut5a toward a soluble substrate, as evidenced by the comparable *p*NPB activities of the two enzymes (Figure 6c). Although enzyme adsorption to the polymer surface was not directly assessed, the substantially greater release of polyester-derived monomers and the more pronounced reduction in turbidity observed for *Fo*Cut5a-*Tr*CBM1 (Figure 6d, e) suggest that the *Tr*CBM1 module promotes a more effective interaction with Impranil DLN-SD, thereby facilitating access to ester bonds within the poly(ester-urethane) and enhancing depolymerization. This interpretation is further supported by the increased accumulation of degradation products observed in vivo for strain YM4B5105 (Figure 5).

## 4. Discussion

In this study, we report the first application of the cutinase *Fo*Cut5a from *F. oxysporum* for poly(ester-urethane) degradation through heterologous expression and secretion in *Y. lipolytica*. This work expands previous studies employing engineered *Y. lipolytica* strains for polyester degradation, which were based on lipases and focused on non-polyurethane substrates, including poly(ε-caprolactone) (Zdanowska et al., 2026). *Fo*Cut5a was identified as the most promising candidate among four esterases based on the extracellular activity detected in the yeast host (Figure 1b).

To explore how modular architecture influences poly(ester-urethane)-degrading performance, a combinatorial library of 96 constructs was assembled by Golden Gate cloning and integrated site-specifically into the *Y. lipolytica* genome (Figure 2). The use of *Y. lipolytica* as an expression host is advantageous given its established capacity for high-level secretion of heterologous proteins and its ability to metabolize hydrophobic substrates (Fickers et al., 2005). Site-specific integration ensured stable single-copy expression, enabling direct phenotypic comparison across construct variants. Among the best-performing colonies, specific module combinations were enriched (Figure 3b-d), suggesting that efficient polyurethane degradation depends on the coordinated interplay of secretion, linker composition, and substrate-binding domains. This study presents the first combinatorial modular platform for engineering poly(ester-urethane)-active enzymes in a yeast host, coupling library assembly with direct phenotypic selection on polyurethane-containing medium.

LC-MS analysis of culture supernatants from strain YM4B5105, a representative variant selected from the library screening (Figure 3e), revealed the accumulation of adipic acid, neopentyl glycol, and 1,6-hexanediol (Figure 4a, b and Figure S1), consistent with the proposed poly(hexane/neopentyl adipate) polyester structure of Impranil DLN-SD (Biffinger et al., 2015; Fuentes-Jaime et al., 2022; de Witt et al., 2024). The detected products indicate hydrolytic cleavage of ester bonds within the polyester soft segment of the poly(ester-urethane), in agreement with the established preference of cutinases for ester rather than urethane linkages (Schmidt et al., 2017; Magnin et al., 2019). Consistently, no HDI-derived compounds were detected, suggesting that degradation was largely confined to the polyester domain. While degradation products from Impranil DLN-SD have previously been characterized in bacterial systems, this study provides the first qualitative and quantitative characterization of monomer release during polyurethane degradation by an engineered yeast platform.

Whereas adipic acid and neopentyl glycol accumulated throughout the incubation, 1,6-hexanediol decreased after an initial accumulation phase (Figure 4a,b). In the absence of significant biomass formation, this decline likely reflects further metabolism rather than incorporation into biomass. Consistent with the known ability of *Yarrowia lipolytica* to utilize 1,6-hexanediol (Carniel et al., 2023), LC-MS analysis identified 6-hydroxycaproate as a downstream product (Figure S1d), extending to a C6 diol the oxidative metabolism previously described for ethylene glycol and 1,4-butanediol in this yeast (Carniel et al., 2023; Messina et al., 2026). The absence of 6-hydroxycaproate in cell-free reactions (Figure 6e) indicates that its formation results from endogenous *Y. lipolytica* metabolism following the release of 1,6-hexanediol rather than from *Fo*Cut5a activity. To our knowledge, this is the first evidence that *Y. lipolytica* converts polyurethane-derived 1,6-hexanediol into 6-hydroxycaproate during poly(ester-urethane) degradation. Although 6-hydroxycaproate levels subsequently declined, its ultimate fate remains unclear. The decrease suggests further metabolic transformation, but complete conversion to adipic acid is not supported by the metabolite profiles, as adipic acid concentrations remained essentially stable while 6-hydroxycaproate levels decreased (Figure 4a-b).

HPLC analysis of the culture supernatants collected from the four selected strains (Figure 3e) after incubation with Impranil DLN-SD (Figure 5) showed that the empty-backbone control strain also accumulated detectable amounts of adipic acid and 6-hydroxycaproate. These findings suggest that *Y. lipolytica* possesses an intrinsic capacity to contribute to the turnover of polyester-polyurethane-derived degradation products. Nevertheless, expression of *Fo*Cut5a variants enhanced adipic acid release up to 2.5-fold above this baseline, with YM4B5105 (*Fo*Cut5a-*Tr*CBM1) exhibiting the highest levels of both metabolites.

Since adsorption to insoluble polymer surfaces is often a major factor limiting polyester hydrolysis, the results obtained with YM4B5105 suggested that fusion of *Tr*CBM1 could improve the interaction between *Fo*Cut5a and the polyester domains of Impranil DLN-SD. Consistent with this hypothesis, direct comparison of YM4B5105 (SigPep2-*Fo*Cut5a-Linker2-*Tr*CBM1) and the parental strain YM4B615 (SigPep2-*Fo*Cut5a) revealed significantly higher accumulation of adipic acid and 6-hydroxycaproate in the former (Figure 6a, b).

The two purified enzymes displayed comparable activity towards *p*NPB (Figure 6c), indicating that addition of *Tr*CBM1 does not alter the intrinsic catalytic properties of *Fo*Cut5a on a soluble substrate. In contrast, the fusion protein was markedly more active on Impranil DLN-SD, as demonstrated by enhanced turbidity reduction and increased release of adipic acid, 1,6-hexanediol, and neopentyl glycol (Figure 6d,e). A closely related cutinase, *Fo*38Cut5, produced in *A. niger*, was recently shown to efficiently degrade the poly(ester-urethane) dispersion Impranil DLN-SD (Verschoor et al., 2025); however, that study assessed degradation solely through optical clearing and did not characterize the released products. Importantly, the enhanced product release observed in vivo was retained with the purified enzymes, indicating that the effect is intrinsic to the enzyme design rather than a consequence of differences in expression or secretion.

Previous studies have shown that binding-domain fusions can improve the performance of PET-active cutinases (Ribitsch et al., 2013; Espino-Rammer et al., 2013) and polyurethane-active polyamidases, although in the latter case improved degradation of the insoluble substrate was accompanied by reduced activity on a soluble model substrate (Gamerith et al., 2016). In that study, a bacterial polymer-binding module enhanced polyamidase activity on polyester-polyurethane copolymers by up to fourfold, while substantially compromising activity on a soluble model substrate. Here, the same strategy was applied for the first time to a poly(ester-urethane)-degrading cutinase, resulting in a 5.6-fold increase in turbidity reduction and 76- and 220-fold higher release of 1,6-hexanediol and adipic acid, respectively, with no detectable change in activity on *p*NPB.

Our data support the hypothesis that *Tr*CBM1 enhances productive enzyme-substrate interactions. Despite being a cellulose-binding module, *Tr*CBM1 is unlikely to act through substrate-specific recognition, as family 1 CBMs bind hydrophobic surface features rather than specific carbohydrate motifs (Lehtiö et al., 2003) . Consistent with this mechanism, fusion of the *T. reesei* CBM1 to a bacterial cutinase enhanced PET hydrolysis 3.8-fold while leaving catalytic activity on a soluble substrate unchanged (Ribitsch et al., 2013). In the case of Impranil DLN-SD, whose colloidal particles provide an extensive hydrophobic surface area, increased residence time at the substrate interface would be expected to increase the proportion of enzyme molecules productively engaged in ester-bond hydrolysis.

## 5. Conclusions

In this study, we demonstrated that the fungal cutinase *Fo*Cut5a, when secreted by *Y. lipolytica*, depolymerizes the polyester segment of the poly(ester-urethane) Impranil DLN-SD, releasing its constituent monomers adipic acid, 1,6-hexanediol, and neopentyl glycol. Fusion of *Fo*Cut5a with the carbohydrate-binding module *Tr*CBM1 significantly enhanced ester-bond cleavage, increasing the release of 1,6-hexanediol and adipic acid relative to the unfused enzyme, without affecting activity on a soluble substrate. Metabolic profiling further revealed the formation of 6-hydroxycaproate, providing evidence that *Y. lipolytica* oxidizes the 1,6-hexanediol released during poly(ester-urethane) degradation. Collectively, these findings establish *Y. lipolytica* as a promising platform for poly(ester-urethane) bioconversion, coupling extracellular depolymerization, to which the endogenous secretome also contributes, with metabolism of the released monomers.

Future work should focus on direct assessment of polymer binding, on extending hydrolytic activity to the urethane bonds of the hard segment, and on enhancing the metabolic assimilation of depolymerization products. This step is essential for poly(ester-urethane) upcycling, as recovery of polymer-derived carbon ultimately depends on its incorporation into host metabolism and subsequent conversion into value-added products. The oxidation of 1,6-hexanediol observed here provides an initial indication that *Y. lipolytica* possesses part of this metabolic potential.

## Acknowledgements

The authors thank Covestro Deutschland AG (Germany) for kindly providing Impranil^®^ DLN-SD. This work was supported by ELIXIR, the European research infrastructure for life science data.

## Author contributions

**Eugenia Messina**: Writing - original draft, Investigation, Methodology, Data curation, Formal analysis; Conceptualization, Visualization. **Markella Papi**: Investigation, Methodology, Formal analysis, Conceptualization. **Serena Barile**: Writing - review & editing, Investigation, Methodology, Data curation, Formal analysis, Visualization. **Rosa De Rosa**: Investigation, Formal Analysis. **Pasquale Scarcia**: Investigation, Methodology. **Antonino Biundo**: Writing - review & editing. **Isabella Pisano**: Writing – review & editing. **Evangelos Topakas**: Writing - review & editing, Project administration. **Gennaro Agrimi**: Writing - original draft, Validation, Supervision, Project administration, Funding acquisition, Methodology, Conceptualization.

## Data availability

All data generated or analyzed during this study are included in this published article and its supplementary information files. Data supporting this study are openly available from Zenodo at https://doi.org/10.5281/zenodo.23321796.

