## Supplementary material for "Synthetic biology-guided modular engineering of the secreted cutinase *Fo*Cut5a with a carbohydrate-binding module improves poly(ester-urethane) depolymerization in *Yarrowia lipolytica*"

**Table S1.** List of primers used in this study

| Name | Sequence (5'-3') | Application |
|---|---|---|
| 1109_For | GCATCGTCTCATCGGGGTCTCATGCC | PCR amplification of the synthetic *Fo*Cut5a gene to introduce BsaI sites for GGA |
| 1179_Rev | CAATGGTCTCTTAGATTATCCAGCAGCCCCGG CAGCATC | PCR amplification of the synthetic *Fo*Cut5a gene to introduce a stop codon and BsaI sites for GGA |
| pTEF For | GTGGTTGGGACTTTAGCCAAGG | Sequencing of the combinatorial libraries and verification of integration at the F11 locus |
| tLIP2 Rev | GTAGATAGTTGAGGTAGAAGTTGTAAAGAGTG | Colony PCR and sequencing of the combinatorial libraries |
| IntF11 For | CGCATAGTTTCTGTCTCCCC | Verification of integration in F11 locus |
| IntF11 Rev | TCAAGTTCTTGCACTCTGTTG | Verification of integration in F11 locus |

| | | |
|---|---|---|
| Barcode_T_For | CTAATTGGCTCTACCGATAGTCTACACCT | Colony PCR and sequencing of the combinatorial libraries |
| Barcode 2 Rev | GGCGTAAATGTTTTCACCAACTTCTTACACC | Sequencing of the combinatorial libraries |
| 1005_1F | AATGAAATCTCTATTGCTGTCGCTGCTGGCGG TCCCGGCCAC | Oligonucleotides forward for Sigpep 1 |
| 1006_1R | GGCAGTGGCCGGGACCGCCAGCAGCGACAG CAATAGAGATTT | Oligonucleotides reverse for Sigpep 1 |
| 1007_2F | AATGAAGCTTTCCACCATCCTTTTCACAGCCT GCGCTACCCTGGCTGC | Oligonucleotides forward for Sigpep 2 |
| 1008_2R | GGCAGCAGCCAGGGTAGCGCAGGCTGTGAAA AGGATGGTGGAAAGCTT | Oligonucleotides reverse for Sigpep 2 |
| 1009_3F | AATGAAGGTGCTCGCCCTGCTGGTTACTGTCT GCTTTTCCGTTGCCTC | Oligonucleotides forward for Sigpep 3 |
| 1010_3R | GGCAGAGGCAACGGAAAAGCAGACAGTAACC AGCAGGGCGAGCACCTT | Oligonucleotides reverse for Sigpep 3 |
| 1011_4F | AATGAAGCTCGCTACCGCCTTTACTATTCTCAC GGCCGTTCT | Oligonucleotides forward for Sigpep 4 |
| 1012_4R | GGCAAGAACGGCCGTGAGAATAGTAAAGGCG GTAGCGAGCTT | Oligonucleotides reverse for Sigpep 4 |
| 1025_L1_F | ATCACGTCTCATCGGGGTCTCAAGGCCCTCCT GGAGG | Linker 1 amplification forward GGA |
| 1026_L1_R | TTCGCGTCTCTGGTCGGTCTCACAGGGCTGG ACCG | Linker 1 amplification reverse GGA |
| 1027_L2_F | ATCACGTCTCATCGGGGTCTCAAGGCACTGGT ATCCGAG | Linker 2 amplification forward GGA |
| 1028_L2R | TTCGCGTCTCTGGTCGGTCTCACAGGGCCGA GCCGGTG | Linker 2 amplification reverse GGA |
| 1029_L3_F | ATCACGTCTCATCGGGGTCTCAAGGCACCATC CCAGG | Linker 3 amplification forward GGA |
| 1030_L3_R | TTCGCGTCTCTGGTCGGTCTCACAGGGCAAT GGCTGCAG | Linker 3 amplification reverse GGA |
| 1031_L4_F | ATCACGTCTCATCGGGGTCTCAAGGCCCCGT CACTG | Linker 4 amplification forward GGA |
| 1032_L4_R | TTCGCGTCTCTGGTCGGTCTCACAGGGCTGC TGTGG | Linker 4 amplification reverse GGA |

**Table S2.** List of plasmids used in this study

| **Plasmid ID** | **Description** | **Purpose** | **Reference** |
|---|---|---|---|
| p-1553 | pF11US1.2 | Destination vector carrying F11 homology arms and URA3 marker | Yuzbashev et al., 2023 |
| p-1084 | pProC-TEF1 | Plasmid carrying TEF1 promoter | Yuzbashev et al., 2023 |
| p-1595 | pTerC-LIP2 | Plasmid carrying LIP2 terminator | Yuzbashev et al., 2023 |
| p-1135 | pCasNA-IntF11 | Expression of Cas9 and gRNA targeting F11 integration site | Yuzbashev et al., 2023 |

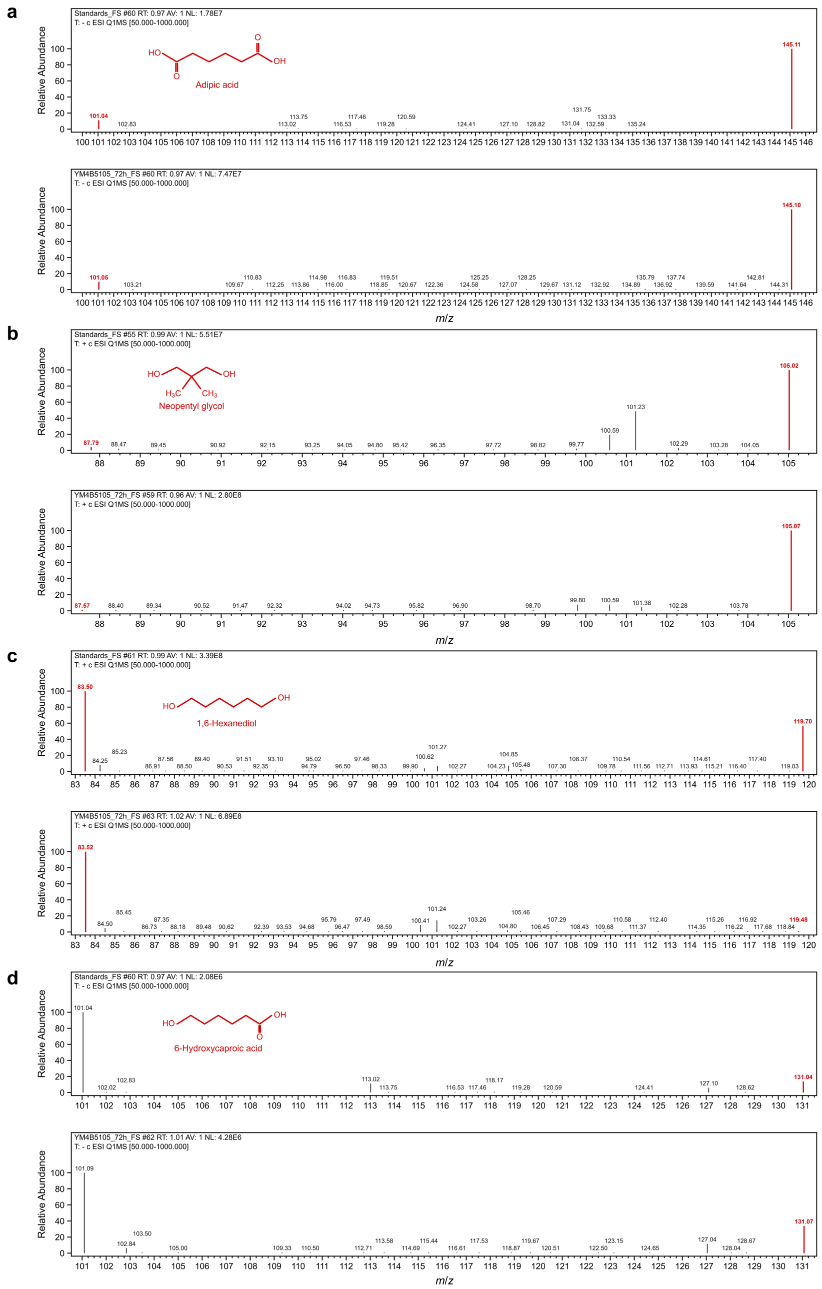


***Figure S1. LC-MS identification of polyurethane monomers released from Impranil DLN-SD in Yarrowia lipolytica culture.*** *Comparison of Q1 full-scan spectra (m/z 50-1000) obtained from authentic standards (upper spectrum of each panel) and the corresponding compounds detected in 72-h culture supernatants of Yarrowia lipolytica YM4B5105 (lower spectrum of each panel): adipic acid (a), neopentyl glycol (b), 1,6-hexanediol (c) and 6-hydroxycaproic acid (d). Adipic acid and 6-hydroxycaproic acid were detected in negative H-ESI mode, showing deprotonated molecular ions at m/z 145.1 ($[M-H]^-$) and m/z 131.0 ($[M-H]^-$), respectively, together with fragment ions at m/z 101.0. In positive H-ESI mode, 1,6-hexanediol and neopentyl glycol yielded protonated molecular ions at m/z 119 ($[M+H]^+$) and m/z 105.0 ($[M+H]^+$), with fragments at m/z 83.5 and m/z 87, respectively. The close correspondence between standard and sample spectra confirms the identity of the detected compounds.*

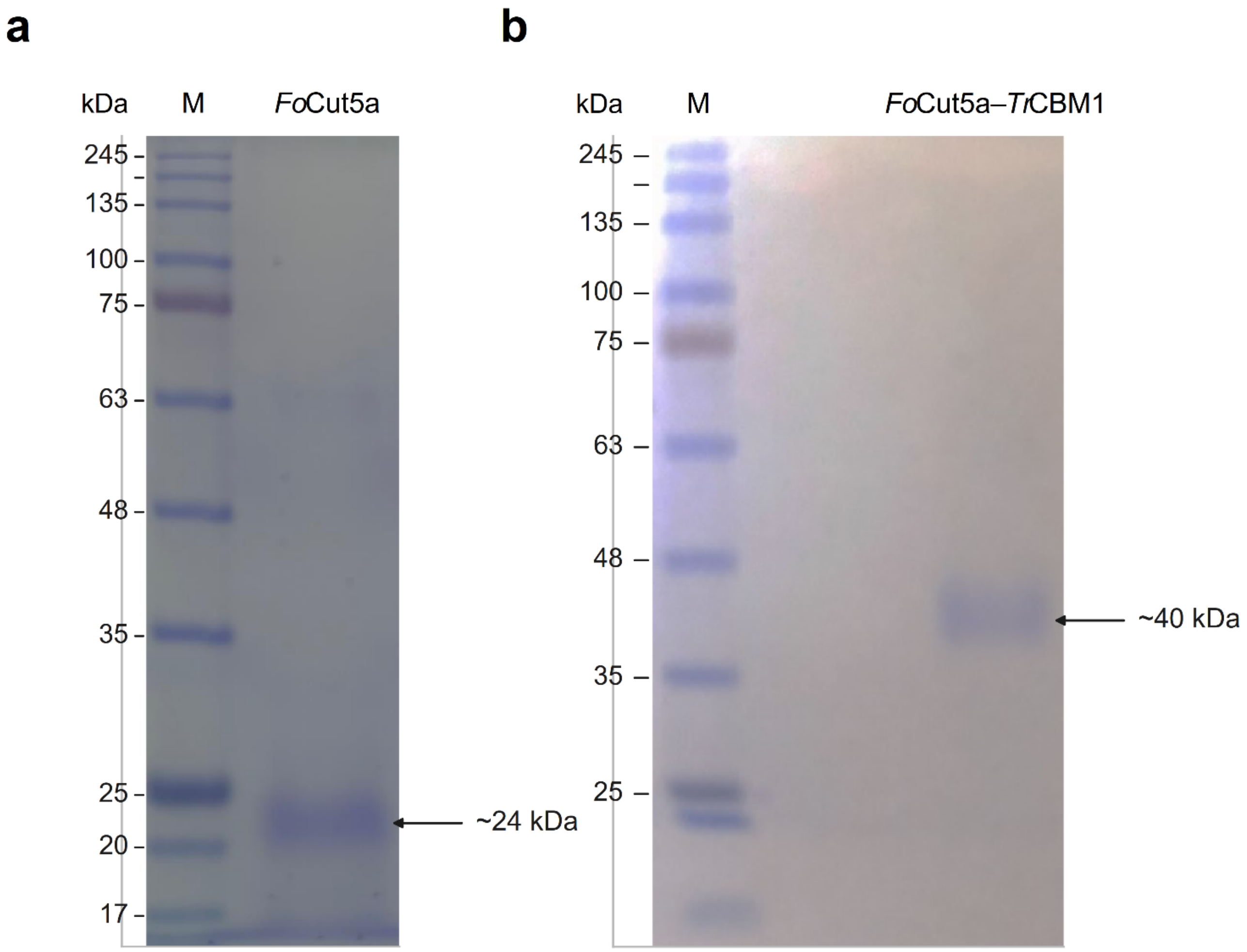


***Figure S2.*** *SDS-PAGE analysis of purified FoCut5a and FoCut5a–TrCBM1.* ***a)*** *FoCut5a purified from strain YM4B615 and* ***(b)*** *FoCut5a-TrCBM1 purified from strain YM4B5105 were separated on 12.5% polyacrylamide gels and visualized by Coomassie Brilliant Blue staining.* ***Lane M****, BlueStar Plus prestained protein marker (NIPPON Genetics); molecular masses are indicated on the left in kDa. Both enzymes migrated as single bands (arrows), with apparent molecular masses of approximately 24 kDa for FoCut5a and 40 kDa for FoCut5a-TrCBM1. The apparent molecular mass of FoCut5a-TrCBM1 was higher than the 35 kDa predicted by the ExPASy ProtParam tool, suggesting glycosylation in the Y. lipolytica expression system. Panels* ***(a)*** *and* ***(b)*** *correspond to independent gels. Images were cropped and subjected to uniform linear adjustments of brightness and contrast across the entire panel; no other image processing was performed.*

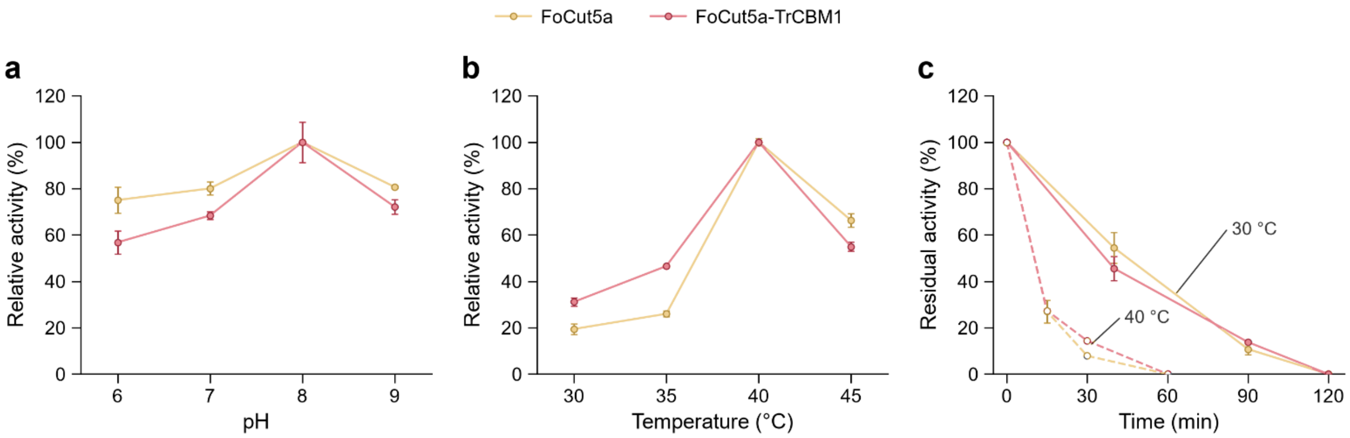


***Figure S3.*** *pH and temperature dependence and thermal stability of FoCut5a and FoCut5a-TrCBM1.* ***(a)*** *Esterase activity, measured as p-nitrophenyl butyrate hydrolysis, at pH 6.0, 7.0, 8.0 and 9.0.* ***(b)*** *Esterase activity at 30, 35, 40 and 45 °C.* ***(c)*** *Residual activity after pre-incubation at 30 °C for 0, 40, 90 and 120 min (solid lines, filled symbols) or at 40 °C for 0, 15, 30 and 60 min (dashed lines, open symbols). Gold symbols, FoCut5a; red symbols, FoCut5a-TrCBM1. In all three panels, symbols are the mean of three technical replicates and error bars are the standard error of the mean (n = 3 in every condition). In panels* ***(a)*** *and* ***(b)****, the activity of each enzyme is expressed as a percentage of its own activity at its optimum condition (pH 8.0 and 40 °C, respectively), which was set to 100%. In panel* ***(c)****, residual activity is expressed relative to the activity measured before pre-incubation (t = 0), which was used as the normalization reference. No residual activity was detected after 120 min at 30 °C or after 60 min at 40 °C. The profiles shown in panels* ***(a)*** *and* ***(b)*** *were not statistically analyzed. In panel* ***(c)****, residual activity was analyzed by two-way ANOVA with incubation time and enzyme as factors, performed separately for each pre-incubation temperature. Incubation time had a significant effect (P = 2 × $10^{-15}$ at 30 °C and P = 2 × $10^{-19}$ at 40 °C), whereas neither enzyme identity (P = 0.51 and 0.19) nor the interaction between the two factors (P = 0.29 and 0.22) was significant. Pairwise comparisons between enzymes at each time point (Welch's t-test with Holm correction) revealed a significant difference only after 30 min at 40 °C (P = 0.0009). Because the three replicates were technical replicates, these statistical analyses describe measurement variability and do not estimate biological variability.*
